# Parallel Emergence and Adaptive Evolution of Multicountry Circulating *Mycobacterium abscessus* Clones

**DOI:** 10.64898/2026.08.02.742357

**Authors:** Chendi Zhu, Mingyu Gan, Yu Zhou, Kelly L Eick, Chase Solomon, Tingting Yang, Anne E Friedland, Jane E. Gross, Melissa B Miller, Kenneth N Olivier, Junhao Zhu, Weimin Li, Qingyun Liu

## Abstract

*Mycobacterium abscessus* (MAB) is a recently emerged bacterial pathogen causing an increasing number of human infections. Despite increasing recognition that different circulating clones contribute disproportionately to MAB population expansion, the global population dynamics and the evolutionary pressures underlying the emergence and spread of MAB clones remain incompletely understood. We analyzed 11,314 publicly available genomes of MAB isolates sampled from 30 countries and identified 38 recently emerged clones that have spread across multiple countries, including the seven previously described Dominant Circulating Clones (DCCs). There was a significant association between the time since the emergence of each circulating clone and its population size, with older clones generally having larger populations. Among the 26 circulating clones of *M. abscessus subsp. abscessus*, 12 carried the macrolide-susceptible *erm(41)* 28C genotype, highlighting clone-level variation in macrolide susceptibility. Genome-wide analyses identified 55 genes that were previously under purifying selection but showed evidence of positive selection during recent clade expansion. Collectively, our findings reveal that the recent global expansion of MAB comprises a broad continuum of newly emerged clones, accompanied by widespread shifts in gene-level selective pressures indicative of ongoing adaptation to new ecological environments.

## Introduction

First recognized as a human pathogen in 1953^1^, *Mycobacterium abscessus* (MAB) has since emerged as an important cause of chronic pulmonary disease in individuals with underlying lung disorders, particularly cystic fibrosis and bronchiectasis, as well as disseminated infections in immunocompromised hosts^2,3^. MAB infections are the second most prevalent nontuberculous mycobacterial infection in humans, and their intrinsic multidrug resistance make them clinically challenging, with limited therapeutic options and average cure rates below 50%^4,5^.

MAB had been regarded as an opportunistic pathogen acquired from environmental reservoirs, but studies of clinical isolates revealed high genomic similarity among strains recovered from patients across distant geographic regions^6,7^. While this observation raised the possibility of person-to-person transmission^8,9^, increasing evidence suggests that this is not the main route of infection^10^; instead, closely related strains may be disseminated through indirect routes, including shared healthcare-associated reservoirs or fomites^11^. A major advance in understanding the MAB population structure came from the identification of seven recently emerged, Dominant Circulating Clones (DCCs) that have disseminated internationally and together account for a large fraction of global, clinical MAB isolates^12^. Despite the predominance of the DCCs, genomic studies have consistently found that 35–50% of clinical isolates fall outside of the seven defined DCCs and are genetically diverse^13–15^, sporadic strains thought to reflect independent environmental acquisition^16^. However, it is not clear whether the non-DCC population consists primarily of unrelated sporadic isolates or includes additional emerging clones that have not yet been recognized. Resolving this question is essential for defining the contemporary population structure of MAB and determining whether ongoing global dissemination is driven exclusively by the seven, defined DCCs or also by other, newly emerging clones.

The emergence and expansion of the DCCs coincided with the advent of the antibiotic era and longer survival of individuals with cystic fibrosis and other immunocompromising conditions -- those most susceptible to MAB infections^16,17^. It is therefore suspected that the recent diversification and international spread of DCCs could be associated with a genomic selection for traits that enhance persistence in human-associated environments, colonization of susceptible hosts or dissemination through indirect transmission routes^17,18^. Parallel evolving DCC lineages could share recurrent mutations or other evolutionary signatures, and identifying these signatures may reveal how an environmental mycobacterium adapts to human-associated niches and the evolutionary steps leading to wide dissemination of the emerging clones.

In this study, by analyzing 11,314 publicly available MAB genomes from 30 countries, we uncovered extensive parallel emergence of recently expanded clades, reconstructed their cross-country dissemination, and characterized the dynamics of clade expansion. We developed an evolutionary path–based genotyping framework to distinguish the different clades and identified genome-wide shifts in selection pressures associated with recent human-associated expansion. This work reveals a dynamic global population structure of MAB shaped by asymmetric dispersal, temporal hierarchy in expansion success, and adaptive evolution consistent with ecological transition to human-associated environments.

## Results

### Parallel emergence of recently expanded eDCCs in the MAB population

To characterize the genetic diversity in the global MAB population, we searched the NCBI SRA database and retained 11,314 MAB isolates from 30 different countries for downstream analyses (Fig. 1A, Table S1). Based on average nucleotide identity (ANI), the MAB strains were divided into the three MAB subspecies: subsp. *abscessus* (8,359 isolates), subsp. *massiliense* (2,449 isolates), and subsp. *bolletii* (506 isolates). Most of the strains were isolated from pulmonary sources (86.7%), predominantly from people with CF (78.5%) (Fig. S1A-C). Subspecies-specific phylogenetic trees showed that the MAB population is deeply diverged but includes numerous “comb-like” recently emerged clades, including the seven previously characterized DCC clades (DCC1 - DCC7) (Fig. 1B-D, Fig. S1D-E). In addition to these seven DCCs, we identified another 31 emerging DCCs (eDCCs) meeting the following criteria: i) clade size ≥ 5; ii) average genetic distance between isolates within the cluster ≤ 25 SNPs^12^; iii) ≥ 300 clade-defining SNPs with no intermediate branches; iv) bootstrap value of the clade ≥ 0.95; v) country distribution ≥ 2.

**Figure 1.**
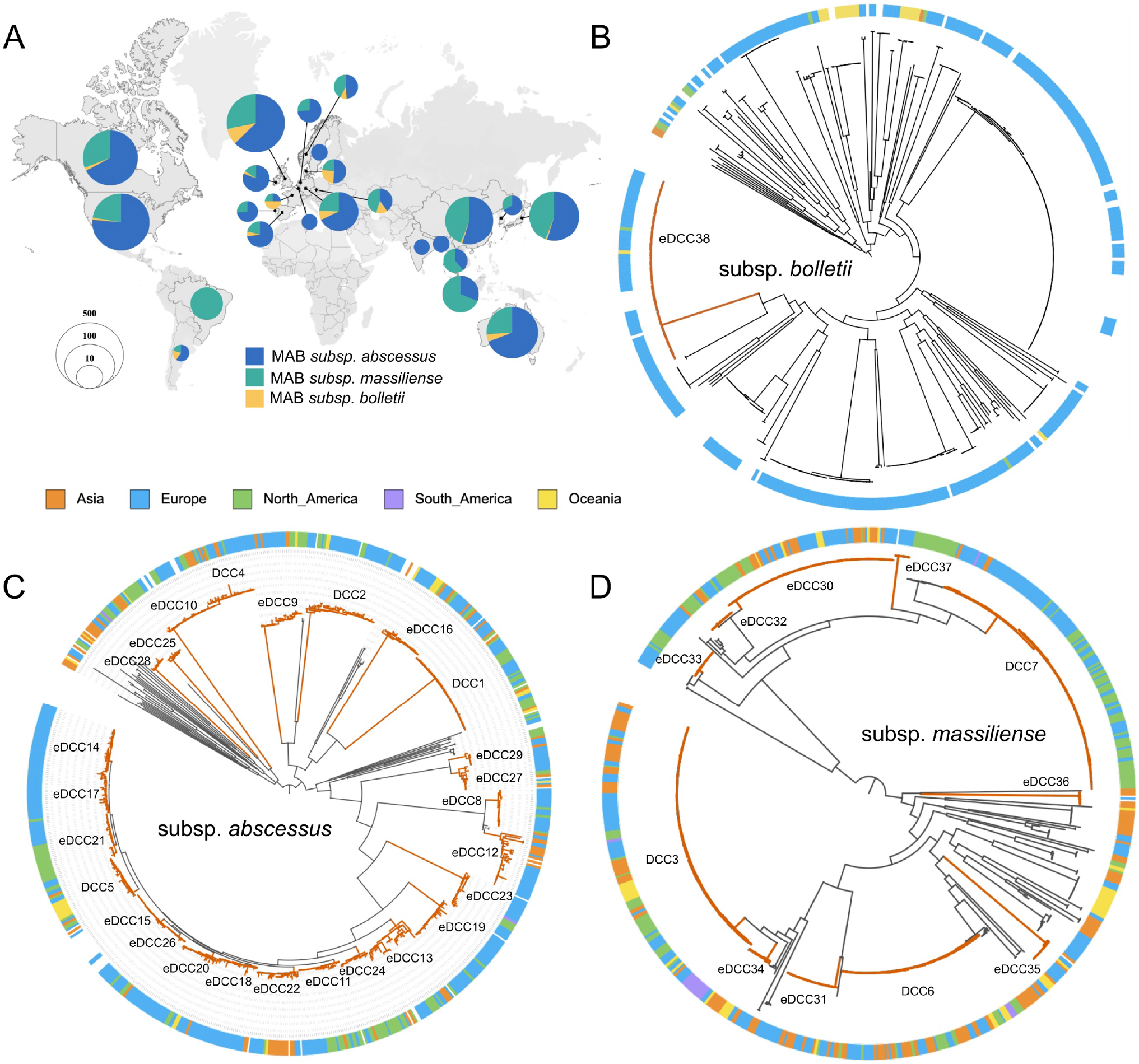
Parallel emergence of circulating clones with cross-country dissemination. (**A**) Geographic distribution of MAB genomes included in this study. Only countries represented by more than three isolates are shown. Pie charts show the relative proportions of the three subspecies at each sampling location: MAB subsp. *abscessus* (blue), subsp*. massiliense* (green), and subsp*. bolletii* (yellow). Circle size indicates the number of isolates sampled. (**B–D**) Circular phylogenies of MAB subsp*. bolletii* (**B**), subsp*. abscessus* (**C**), and subsp*. massiliense* (**D**). Orange branches indicate expanded circulating clades identified in this study. The outer ring denotes the sampling continent of each isolate. To improve visualization of eDCC internal structure, representative subsets of DCC1–3 were included in the main trees; complete trees are shown in Supplementary Figure 1.

The many small, recently emerged clades with multi-country distribution suggested that the number of eDCCs had not yet reached saturation. To test this, we performed a subsampling analysis (similar to rarefaction, Methods) and found that even at the upper limit of sample inclusion, the number of eDCCs continued to increase without reaching a plateau (Fig. S1F), while the proportion of strains not belonging to any DCC decreased (Fig. S1G). In our collection, DCC1–7 accounted for 52.6%(5,952/11,314) of sampled isolates, while the eDCCs accounted for an additional 16.8%(1,906/11,314). Together, these findings demonstrate that the global MAB population is comprised of numerous, independently emerged clades with cross-country distributions, and that the inclusion of more MAB isolates will likely identify additional emerging lineages.

### Asymmetric intercontinental dispersal of DCCs and eDCCs

To investigate the global spread of different MAB clades, we reconstructed the geographic origins of DCCs and eDCCs with Bayesian based ancestral state inference (Methods). Using DCC1 and DCC3 as representative examples (Fig. 2A-C), phylogeographic reconstruction of both clades showed clear signatures of international dissemination following their initial expansion (Fig. 2B, C). DCC1 originally expanded in the United States (posterior probability: 96.1%), followed by secondary expansions in the United Kingdom, China, and Australia (Fig. 2C). DCC3 likely originated in the United Kingdom (posterior probability: 79.35%), but also showed secondary expansions in Brazil, China, and the United States (Fig. 2B). To control for potential sampling bias arising from countries that are overrepresented in our strain collection, we repeated the phylogeographic reconstruction using subsampled datasets with balanced country representation (Methods) and found that the inferred geographic origin patterns persisted (Fig. S2A-F).

**Figure 2.**
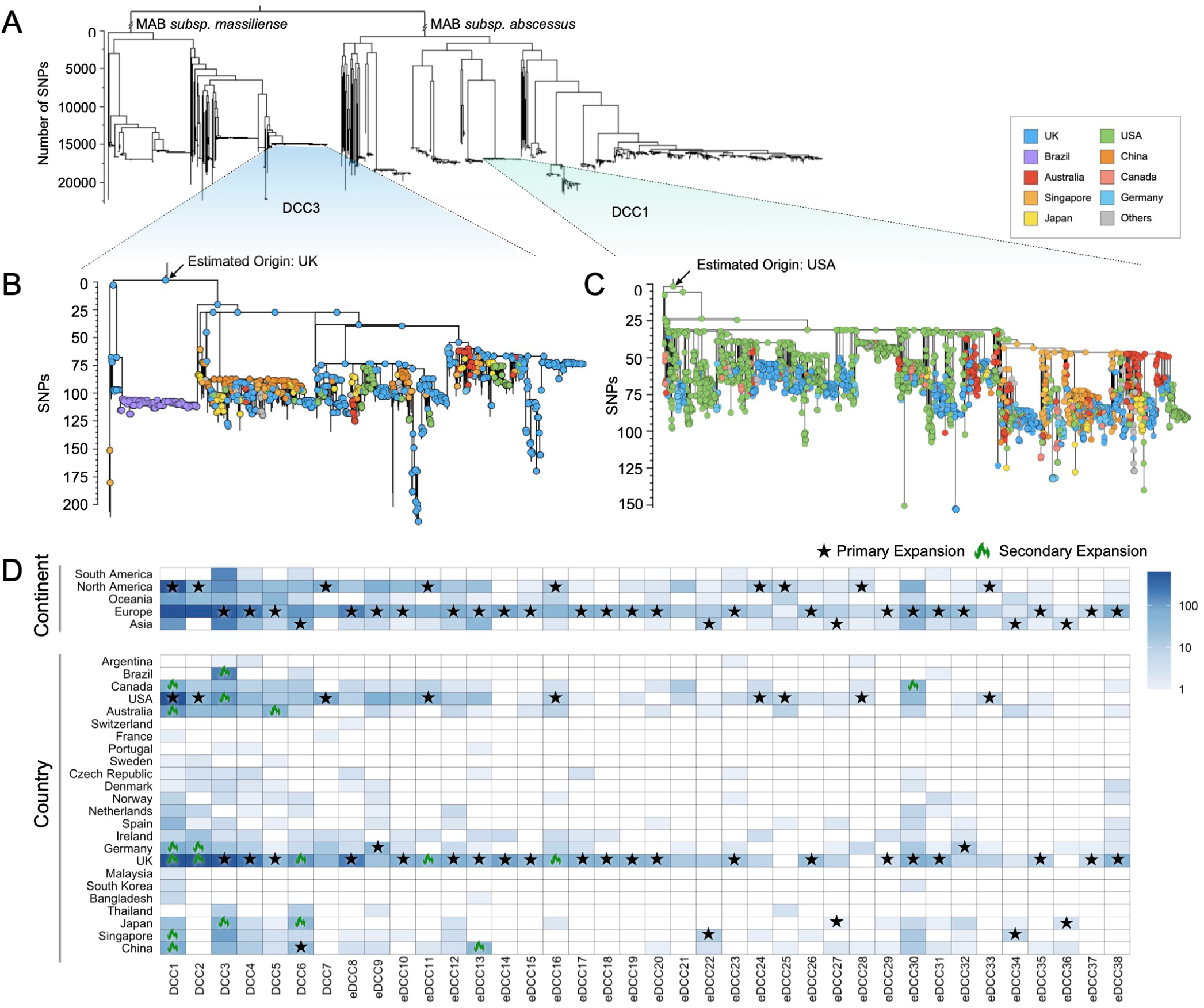
DCCs and eDCCs show asymmetric global dissemination. (**A**) Phylogenetic context of MAB subsp. *massiliense* and *abscessus*, highlighting DCC3 and DCC1 for detailed analysis. (**B–C**) Zoomed-in phylogenies of DCC3 (**B**) and DCC1 (**C**) showing fine-scale population structure and geographic distribution of isolates. Tip colors indicate country of sampling, and arrows mark the inferred geographic origin of each clade. (**D**) Global distribution of DCCs and eDCCs across continents and countries. Only countries contributing more than three isolates were included in the cross-country analysis (n = 24). Color intensity indicates the number of isolates sampled from each region or country. Stars indicate inferred primary expansion events, and flame symbols indicate inferred secondary expansion events.

Cross-country transition events within the other recently expanded clades were inferred from the multi-country origin of isolates within the clades (Methods) (Fig. 2D). Most of the MAB isolates from Asia were phylogenetically nested within clades inferred to have originated in Europe or North America (Fig. 2B,C, Fig. S3). However, five clades—DCC6, eDCC22, eDCC27, eDCC34, and eDCC36—appear to have originated in Asia with limited transmission to Europe and North America (Fig. S3). Together, these observations show that the recent expansion of MAB clades was characterized by an asymmetric, hub-structured pattern of global dispersal.

### eDCCs represent successive waves of recent clonal expansion

We next sought to estimate the time of origin and subsequent expansion dynamics of DCCs and eDCCs. Using a previously defined MAB substitution rate (8.76×10^-8^–2.41×10^-7^ substitutions per site per year)^16^, we inferred the emergence time of the most recent common ancestor for each clade. The majority of clades appear to have emerged within the last century, except for DCC1–3, which originated in the late 19th century (Fig. 3A, Fig. S4A-N). We then assessed the onset of expansion, based on the earliest time at which a clade’s relative genetic diversity increased by more than tenfold compared with the diversity at the phylogenetic root ^16^. Although there was a wide variation in the inferred origin times of the different clades, the expansion of all clades occurred more recently. The expansion of DCC1–3 began in 1961, 1953, and 1953, respectively, while the expansion of the other DCCs and all eDCCs began between 1980 and 2017 (Fig. 3A,B). DCC1–3 exhibited markedly larger and more sustained population growth, while DCC4–7 displayed expansion trajectories comparable to many of the eDCCs (Fig. 3C).

**Figure 3.**
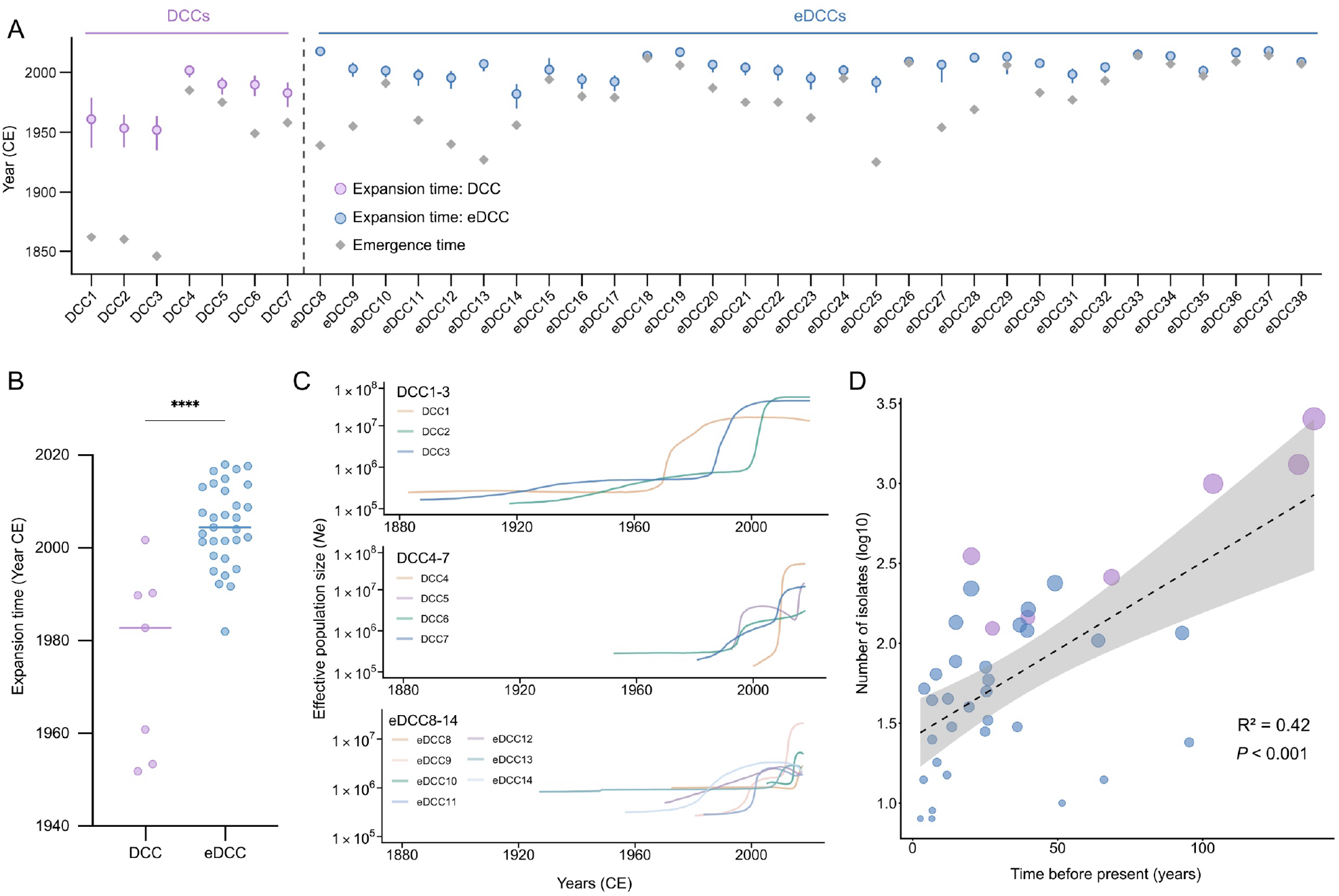
Population dynamics of DCCs and eDCCs. (**A**) Estimated emergence and expansion times of DCCs and eDCCs. Circles indicate estimated expansion times, gray diamonds indicate estimated emergence times, and error bars show 95% confidence intervals. (**B**) Comparison of estimated expansion onset between DCCs and eDCCs (****, *P* < 0.001). (**C**) Bayesian skyline estimates of effective population size (Ne) over time for representative DCCs and eDCCs, grouped as DCC1–3, DCC4–7, and eDCC8–14. (**D**) Relationship between clade age and clade size. Each point represents a DCC or eDCC, with clade size expressed as the log10-transformed number of isolates. The dashed line shows the fitted linear regression, and the shaded region indicates the 95% confidence interval.

All clades, however, showed a significant positive correlation between the time since their origin and their current population size (*R*^2^=0.42*, P*<0.001; Fig. 3D). This correlation persisted when tested with different species-specific substitution rates, with a recently proposed clade-specific mutation rate^19^ (*R*^2^=0.43*, P*<0.001, Fig. S5A) or when the analysis was restricted to strains from individual countries (Fig. S5B-D). Clades that emerged earlier tend to have expanded into larger contemporary population sizes, while more recently emerged clades remain in a stage of active population growth.

### DCC- and eDCC-specific macrolide resistance genotypes and clade genotyping

The MAB clades showed extensive genetic diversity and heterogeneous population dynamics, but we reasoned that using phylogenetics to discriminate the different clades might be valuable for epidemiologic surveillance, clinical diagnostics and patient management. For example, within subsp. *abscessus*, we found 15 clades carrying the 28T sequevar of *erm(41),* which is associated with inducible resistance to macrolides, and 12 clades carrying the 28C sequevar that confers macrolide susceptibility (Table S2, Fig. S6). We therefore sought to establish an evolutionary path–based genotyping scheme to classify the DCCs and eDCCs.

We reconstructed the ancestral sequence of each DCC and eDCC and identified mutations accumulated along the evolutionary path from the ancestral node to the clade ancestor, yielding 478–21,768 clade-defining SNPs per clade (Fig. 4A, Table S3). Comparison of clade-defining SNPs across clades revealed extensive mutation sharing and substantial cross-clade matching (Fig. S7A,B), primarily due to recombination events accumulated during the long-term diversification of the MAB population (Fig. S7C). We therefore limited the barcoding SNP sets by considering only SNPs present in ≥95% of isolates within the target clade, and in <5% of isolates outside the clade. This filtering reduced the clade-defining barcode sets to 8–9,061 SNPs per clade. Using these refined SNP sets, DCCs and eDCCs were successfully identified, with isolates carrying 95.4%–100% of the clade-defining SNPs defining their corresponding clades (Fig. 4B). The SNPs sets distinguished DCC/eDCC isolates from non-DCC isolates, all of which carried <75% SNP of the clade-defining SNPs for any DCC/eDCC clade (Fig. 4C). Thus, the barcode SNPs can accurately identify the different MAB clades, providing a scalable tool for genomic surveillance, epidemiological tracking and antibiotic selection.

**Figure 4.**
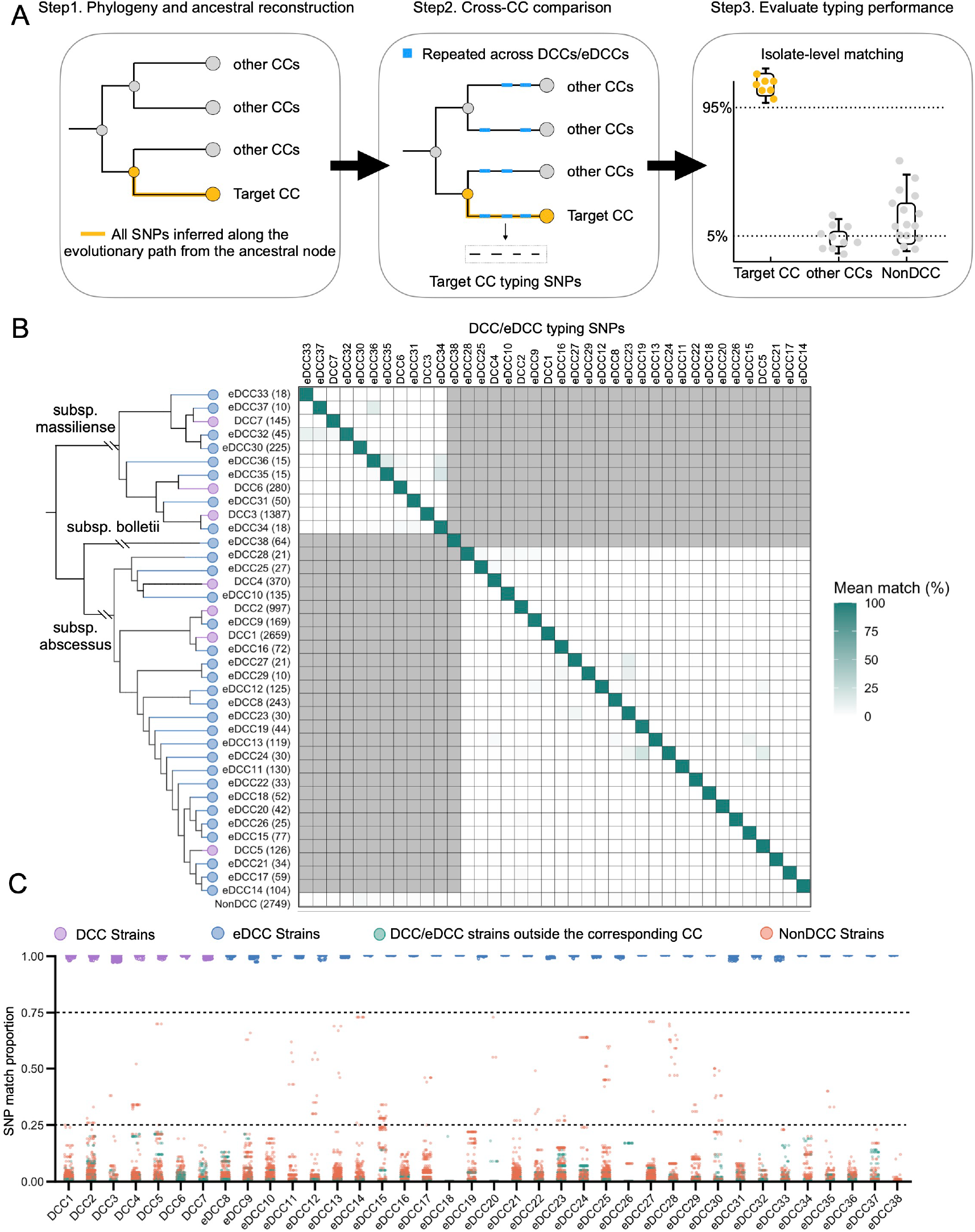
Clade-defining SNPs distinguish DCCs and eDCCs. (**A**) Schematic overview of the strategy used to identify DCC/eDCC-defining SNPs. SNPs arising along the evolutionary path from the ancestral node to a target clade were identified, compared with all other DCCs/eDCCs, and retained as typing SNPs if they were specific to the target clade. (**B**) Heatmap showing the specificity of selected DCC/eDCC-defining SNPs for assigning isolates to DCCs and eDCCs. Values indicate the mean percentage of SNPs matched by each isolate group (rows) to each DCC/eDCC-specific SNP set (columns). Gray shading indicates comparisons across subspecies. (**C**) Isolate-level performance of selected DCC/eDCC-defining SNPs. Purple and blue points indicate isolates from the corresponding DCCs and eDCCs, respectively; orange points indicate non-DCC/non-eDCC isolates, and green points indicate isolates from non-corresponding DCCs or eDCCs. Isolates from the target clade showed near-complete SNP matching, whereas isolates outside the target clade showed substantially lower matching proportions.

### Parallel adaptive evolution during recent DCC and eDCC expansion

The recent expansion of MAB clades in isolates from human infections suggests ongoing adaptation to human-associated environments. We performed gene-level *pN/pS* analyses to identify loci under positive selection (Methods), focusing on mutations that accumulated after clade expansion. This analysis identified 55 genes under positive selection, including 14 previously reported targets and 41 newly identified candidates (Fig. 5A, Table S4). We also observed a higher-than-average mutation frequency in a subset of intergenic regions that included the promoters for *espR* and *whiB1* (Fig. 5A, Fig. S8A), and there were also positive selection signals in nearby protein coding regions. This suggests that adaptive evolution in MAB may include mutations in regulatory elements that modulate gene expression. Analysis of individual circulating clones revealed that the mutations in many of these genes occurred independently across multiple DCCs and eDCCs, and were generally nonsynonymous changes (Fig. S8B). Taken together, these findings indicate parallel selection across circulating clones, with DCCs and eDCCs subject to similar selective pressures.

**Figure 5.**
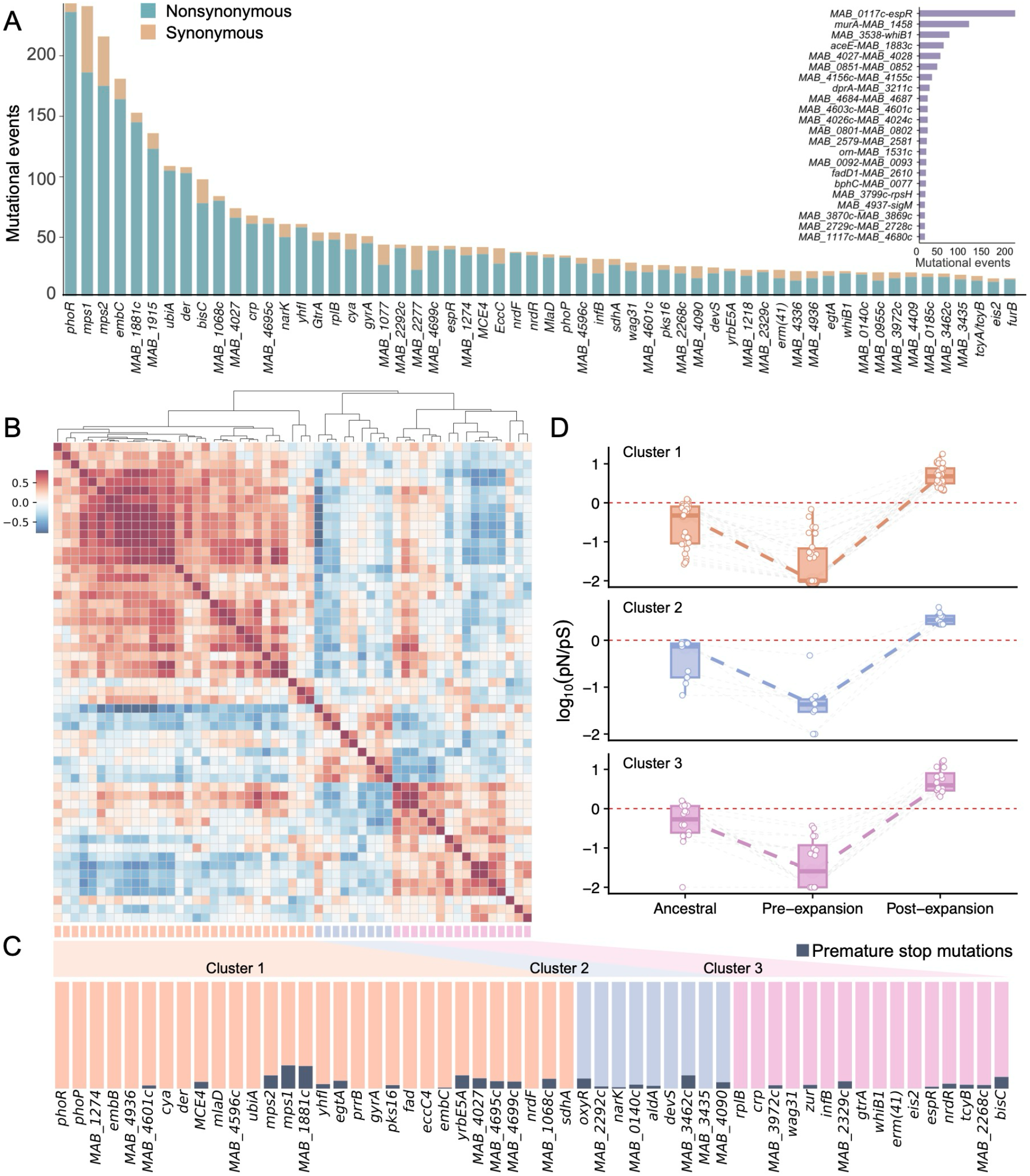
Positively selected genes form co-expression modules associated with MAB adaptation. (**A**) 55 ranked by the numbers of nonsynonymous and synonymous mutations accumulated after clade expansion. The inset shows mutational counts in recurrently mutated intergenic regions (IGRs). (**B**) Co-expression patterns among the 55 positively selected genes. The heatmap represents pairwise Pearson correlation coefficients, with hierarchical clustering grouping genes according to their co-expression patterns. (**C**) Assignment of the positively selected genes to three co-expression modules. Dark blue indicates the proportion of premature stop mutations. (**D**) Changes in selective pressure across evolutionary stages for genes in each co-expression cluster. Boxplots summarize the distributions of log10(pN/pS) during the ancestral (within-subspecies diversification), pre-expansion (DCC/eDCC defining), and post-expansion stages. The horizontal dashed line indicates log10(pN/pS) = 0.

We reasoned that genes mutated under positive selection may not act in isolation, but instead may work together in functional modules associated with biological processes undergoing adaptive remodeling. Our initial assessment, using existing pathway annotations to determine whether these 55 genes were functionally interconnected, yielded only limited results, likely because a substantial fraction of the 55 genes are annotated as having unknown functions. We therefore took an annotation-agnostic approach, examining 129 publicly available MAB transcriptomic datasets to see if these genes were transcriptionally associated^20^. We found that these 55 genes clustered into three co-expression modules (Pearson’s correlation coefficient > 0.5; Fig. 5B-C). Each of the three clusters contains two or more key regulators: *phoP/R* in Cluster 1, *devS* and *oxyR* in Cluster 2, and *crp*, *zur*, *espR* and *whiB1* in Cluster 3. The three co-expression clusters were associated with distinct biological processes: Cluster 1 with cell envelope biosynthesis (*ubiA, embC,* and *embB*) and lipid metabolism (*mce4, mlaD*, *yrbE5A, mps1, mps2, pks16*, and *eccCa*); Cluster 2 with redox homeostasis and energy metabolism (*oxyR, devS, aldA, narK, nrdF, nrdR,* and *sdfA*); and Cluster 3 with cell growth and division (*rplB, crp, cya, wag31, infB*, and *bisC*) (Fig. 5C, Table S4). There were also two genes associated with antibiotic resistance: *erm(41)* (inducible macrolide resistance) and *eis2* (aminoglycoside resistance).

Finally, to determine whether the positive selection on these three modules occurred during the recent expansions, we compared selection signals before and after clade expansion (Methods). We found that genes in all three modules were under purifying selection during long-term evolution, as reflected by mutations that accumulated after subspecies divergence but before the emergence of DCCs/eDCCs, and subsequently shifted toward positive selection during DCC/eDCC expansion (Fig. 5D, Table S4). This selective pressure shift suggests that genes previously maintained under evolutionary constraints became recurrent targets for mutations during recent clonal expansions, presumably reflecting adaptation to human-associated environments.

### Expansion-associated adaptive variants disseminate across patients and countries

We next examined whether adaptive variants in genes under positive selection that were acquired during clade expansion were retained in subsequent human isolates, suggesting continued propagation in human populations. We identified 1,314 potential propagation events (Fig. 6A–B, Table S5), in which the acquired mutations were present in isolates from different patients within a genomic cluster. Among these, 211 events involved isolates from different countries, suggesting propagation across national borders (Fig. 6A–B, Table S5). For example, in the Brazilian outbreak clade (DCC3) ^21^, the *embC* L78F and *MAB_4691c* A4275V mutations, acquired before expansion, were present in isolates from different geographic regions, showing that the adaptive changes preceded geographic dissemination (Fig. 6B). In addition to these two mutations, the clade subsequently acquired *rrl* A1381G and *MAB_1077* L169P and continued to expand among patients (Fig. 6B). Collectively, these findings indicate that acquired adaptive mutations are maintained across individuals and international borders.

**Figure 6.**
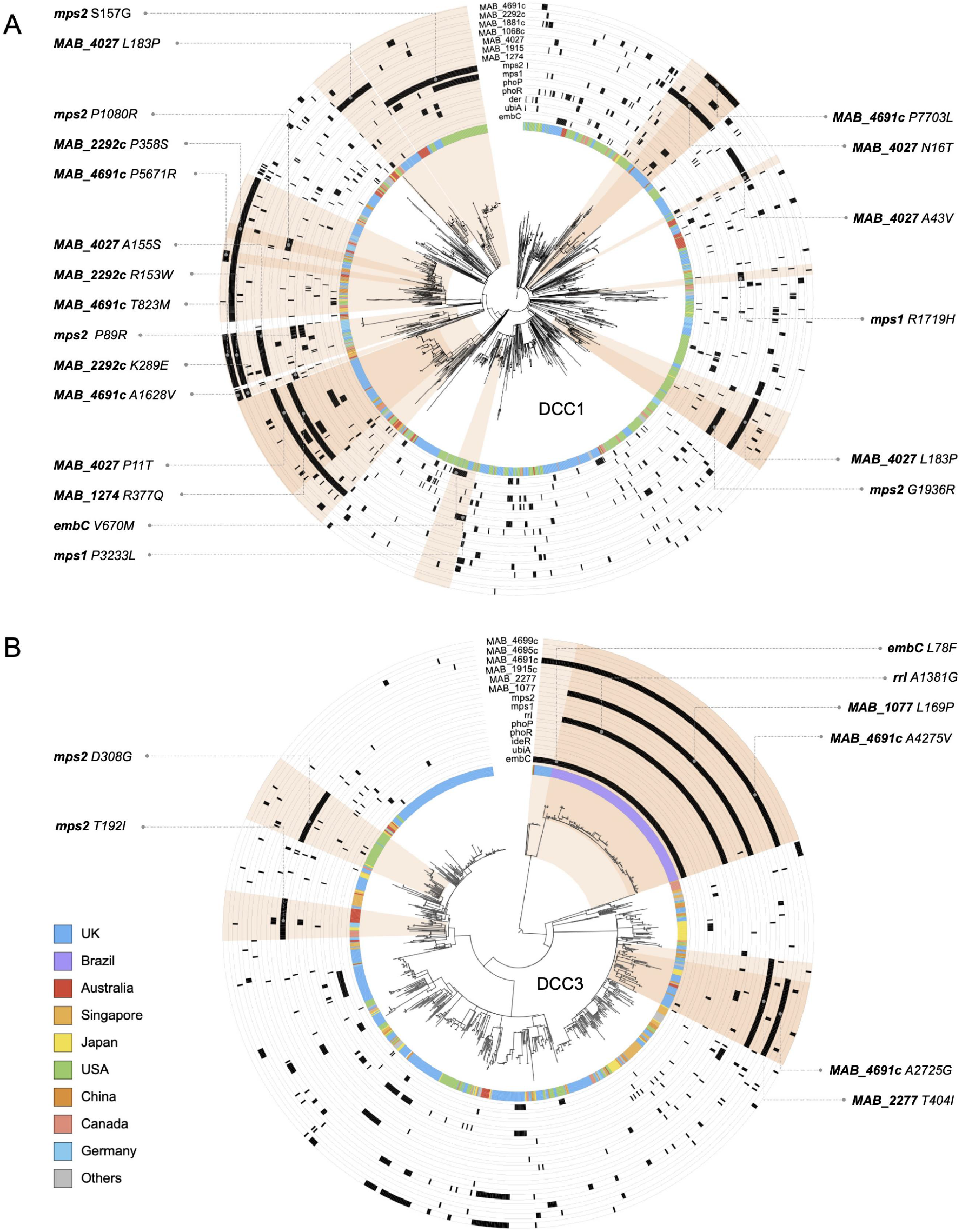
Propagation of mutations in positively selected genes across patients in DCC1 and DCC3. Circular phylogenies of DCC1 (**A**) and DCC3 (**B**) showing the distribution of mutations in genes under positive selection. The inner colored ring indicates country of sampling, and outer black tracks mark the presence of mutations in selected genes. Shaded sectors highlight genomic clusters in which identical mutations were shared among isolates from multiple patients, suggesting onward transmission of strains carrying these mutations. Representative shared mutations are labeled around each tree.

## Discussion

In this study, we show that the majority of global MAB isolates belong to a series of recently emerged clades. We reconstructed the geographic origins and intercontinental dispersal routes of these clades, uncovered asymmetric patterns of global dissemination, and identified 55 genes under positive selection during recent expansion. Together, these findings provide a systematic view of MAB emergence, linking parallel clade expansion and global dispersal with adaptive shifts toward human-associated environments.

The MAB population is structured into a large number of deeply separated phylogenetic clades. Within each clade the strains differ by an average of ≤ 25 SNPs, whereas strains from different clades differ by an average of >10,000 SNPs (2,214 – 33,173) (Fig. S9), highlighting a pattern of discontinuous diversification. However, because the majority (98.3%) of available genomes were isolated from humans, the discontinuous structure may reflect sampling bias toward disease-associated strains, while environmental genotypes, which might bridge these phylogenetic gaps, are underrepresented. Similar host-associated phylogenetic clustering has been reported in pathogens such as *Staphylococcus aureus* ^22^ and *Pseudomonas aeruginosa* ^23^, in which human isolates belong to clades that are phylogenetically separate and distinct from environmental populations. Alternatively, a recent study proposed that this type of broom-like phylogenetic structure arises through genome-wide selective sweeps, in which more adapted clones replace their less adapted closest relatives before re-diversifying into ecologically differentiated populations^24^. Currently, because genomic data from environmental MAB strains is limited, it is impossible to distinguish between these models, and additional environmental MAB genomes are needed to clarify the ecological forces driving the recent MAB expansion.

The recent expansion of MAB has been related to changes in host demographics and clinical practice. The most prominent association is with the increasing lifespan of individuals with CF^16^, who are often prescribed prolonged and intensive antibiotic treatments^25^ that may have created a novel ecological niche favorable to MAB. Previous studies found that the MAB genome contains genes unique to other bacterial species frequently colonizing people with CF, such as *Burkholderia* spp. and *Pseudomonas aeruginosa*, and these genes, including *mgtC* and *plc*, may contribute to the high incidence of MAB infections in people with CF^26,27^. In addition, the acquisition of new genes, such as *dpnM*, through horizontal gene transfer, could play a role in the emergence of DCCs^11^. A systematic analysis of MAB pan-genomes, based on DCCs and eDCCs from human infections, together with additional environmental isolates from diverse sources, may help to identify shared genetic determinants promoting the expansion of these clonal complexes.

The temporal hierarchy in clade expansion suggests that a clade’s population size reflects not only the time since its establishment, but perhaps also its progressive adaptation to human-associated environments. Earlier-emerging clades may have gained access to human transmission networks sooner, allowing cumulative demographic expansion, but their persistence and sustained growth might also be the result of a gradual accumulation of adaptive mutations facilitating survival in antibiotic-exposed or host-associated niches. Consistent with this interpretation, isolates belonging to DCC1–3 were significantly more likely to carry adaptive mutations than isolates from other clonal complexes (63.2% vs. 46.9%, *P* < 0.001; Fig. S10), suggesting that prolonged circulation in humans provides greater opportunities for adaptive evolution. The continued emergence of newer clades suggests that this adaptive process remains ongoing and is not yet saturated.

While we observed an apparent dominance of DCCs 1-3, we can’t assess whether this reflects pathogenic superiority because we don’t have the epidemiological data to assess whether certain clones competitively displace others following introduction into a population, a pattern frequently observed in other bacterial pathogens^22^. Longitudinal surveillance and integrated clinical datasets will be required to determine whether specific clades differ in transmissibility, persistence, or clinical severity. MAB pathogenicity has been studied in preclinical models, such as immunocompromised mice^28^, *Drosophila*^29^, and zebrafish embryos^30^, and it would be intriguing to test whether clades exhibiting distinct epidemiological patterns differ in their ability to infect and persist in these animal models. Also, while subsp. *abscessus* has historically been associated with inducible macrolide resistance via the *erm(41)* 28T sequevar^31^, we found that resistance patterns segregate at the clade level. Thus, clade based genotyping provides higher resolution for clinically relevant traits than subspecies designation alone.

Another major finding of this study was the identification of 55 genes that were subjected to positive selection during recent clade expansion and could therefore be studied for possible roles in drug susceptibility or host-bacteria interactions. We showed that these putative adaptive mutations are not confined to within-host evolution, but can be disseminated across hosts and international borders. However, the frequency of these adaptive mutations in the MAB population remains low. Only five genes (*phoR*, *mps1*, *embC*, *mps2*, and *MAB_1881c*) were mutated in more than 5% of total isolates. The most common, *phoR*, was mutated in just 6.78% of isolates (Table S6), and the remaining genes were mutated in only 0.03–4.67% of isolates, suggesting that adaptation to the lung environment is still at an early stage. Notably, signals of positive selection differed between MAB subspecies. *MAB_1881c*, *MAB_1068c*, *bisC*, and *MAB_2292* were mutated only in subsp. *abscessus* clades, whereas *MAB_1077*, *MAB_4336*, and *MAB_2277* were mutated only in subsp. *massiliense* clades (Fig. 5B, Table S4). This suggests that shared selective pressures during human-associated expansion may act through subspecies-specific genetic targets. Future functional studies will be needed to determine whether these mutations represent alternative evolutionary routes to similar adaptive traits.

Our study has several limitations. First, because our analysis relies on publicly available genomes, the uneven sampling across regions and clinical populations may have caused some expanding MAB clades to be underrepresented or missed entirely. Second, although the genomic signatures we identified are consistent with recent adaptation, functional validation is needed to establish possible mechanistic links between candidate mutations and antibiotic susceptibility, host-associated fitness, or environmental persistence. Third, we did not collect clinical metadata and treatment outcomes, and thus cannot determine whether clade identity or adaptive genotype predicts disease severity, persistence, or treatment response. Finally, we did not analyze selective pressure on the accessory genome because the presence or absence of accessory genes may reflect clonal background, hitchhiking, or lineage-specific gene gain or loss rather than adaptive selection.

In summary, our findings redefine MAB emergence as a broad and ongoing evolutionary process rather than the expansion of a few established dominant clones. By analyzing MAB genomes outside of the seven previously defined DCCs, we identified additional, recently expanded lineages at distinct stages of clonal emergence and global dissemination. The recurrent signals of positive selection associated with these expansions suggest that clonal success is coupled with an evolutionary process of adapting to human-associated niches. This framework provides a basis for tracking how new MAB clones arise, spread, and evolve from environmental opportunists into globally disseminated human-associated pathogens.

## Materials and Methods

### Genome sequences of MAB isolates

We searched PubMed (as of March 1, 2025) to identify articles that published whole-genome sequencing data for *Mycobacterium abscessus* strains, using the keyword “*Mycobacterium abscessus”* or “*Mycobacteroides abscessus*”. Concurrently, we also searched the NCBI SRA database using the keyword "*Mycobacterium abscessus*" or "*Mycobacteroides abscessus* ". We identified 188 BioProjects comprising 11,702 whole-genome sequencing samples of MAB. Following species identification, removal of low-quality or contaminated sequences, and other quality-control procedures, 11,314 nonredundant MAB isolates were retained for downstream analyses (Table S1). These samples were originally collected from 30 countries or regions across Asia, Europe, Oceania, and the Americas (Table S1). The information about longitudinal samples from the same patient was extracted from the BioSample database or the supplementary data from the published papers (Table S1).

### Subspecies identification

We performed *de novo* assembly of all sequencing reads using SPAdes v3.11.1^32^. Specifically, we used "--careful" parameter and set the PHRED quality offset to 33 using the "--phred-offset 33" parameter. These contigs were then compared to reference sequences for each of the three subspecies: subsp. *abscessus* GZ002 (NZ_CP034181.1), subsp. *massiliense* CCUG48898 (NZ_AP014547.1) and subsp. *bolletii* GD91 (NZ_CP065265.1). To identify the subspecies assignment of each MAB strain, we calculated a whole-genome-based average nucleotide identity (gANI) score using *fastANI* (v1.2)^33^. Each MAB strain was then assigned to a subspecies if it had a gANI score of at least 98% compared to the reference strain of that subspecies (Table S1). All 11,314 MAB strains were successfully assigned to a specific subspecies.

### SNP calling

The *Sickle*^34^ tool was used to trim the WGS data, and sequencing reads with a Phred base quality above 20 and read lengths longer than 30 were kept for analysis. *BWA MEM* (v0.7.17)^35^ was used to map sequencing reads using the corresponding subspecies reference sequences as templates. *SAMtools* (v1.3.1)^36^ was used for SNP calling with sequencing depth ≥ 20. Fixed mutations (frequency ≥95%), unfixed mutations (5%≤ frequency <95%), and indels were identified using *VarScan2*^37^ (v2.3.9). SNPs in repetitive regions of the genome (phage sequences, insertions, and mobile genetic elements) were excluded.

### Phylogenetic reconstruction

The SNPs of the MAB isolates were catenated into a single consensus and non-redundant list while nucleotide positions with gaps in more than 5% of the taxa were excluded as possibly due to insertions, deletions, low coverage, or poor mapping quality at those sites. We used *Fasttree*^38^ and the General Time-Reversible (GTR) model of nucleotide substitution with four gamma rate categories to inferred maximum likelihood (ML) phylogenetic trees for subsp.*abscessus* (8,359 isolates), subsp. *massiliense* (2,449 isolates), and subsp. *bolletii* with (506 isolates). To further validate the robustness of the trees, a maximum likelihood (ML) phylogenetic tree was inferred using *IQ-TREE* (v2)^39^, with ultrafast bootstrap supports from 1,000 replications. The best-fit nucleotide substitution model was GTR+I+G, as determined by *ModelFinder*^40^. Phylogeny trees were visualized in *FigTree* (v1.4.4)^41^ or *iTOL*^42^.

### Identification of emerging dominant circulating clones (eDCCs)

Candidate eDCCs were identified from phylogenetic clusters outside the seven previously defined DCCs. To maximize the detection of potential early-stage dominant clones while minimizing false-positive local transmission clusters, we established five criteria for eDCC identification. First, each candidate clade was required to contain isolates from at least five independent patients. This lower threshold was chosen to capture recently emerging clonal complexes before they reached the size of established DCCs while reducing the likelihood of including sporadic transmission events. Second, the mean pairwise genetic distance within each clade was required to be ≤25 SNPs, consistent with the genomic diversity observed in recently expanded MAB clones, ensuring that each candidate represented a genetically coherent lineage^12,16^. Third, each candidate clade had to be separated from its nearest sister lineage by an uninterrupted ancestral branch containing at least 300 ancestral-branch SNPs. This criterion was introduced to distinguish independently evolved lineages from shallow phylogenetic splits or derivatives of existing clonal complexes. Under the estimated substitution rate of 1.8 – 2.2 SNPs per genome per year, this branch length corresponds to approximately 140 – 170 years of independent evolution, substantially exceeding the timescale over which currently recognized DCCs have diversified^16^. Consequently, this threshold minimizes the possibility that candidate eDCCs simply arose through recent acquisition of adaptive variants from established DCCs and instead enriches for independently evolved clonal complexes. Fourth, the ancestral node defining each candidate clade was required to have bootstrap support ≥ 0.95 to ensure phylogenetic robustness. Finally, each candidate clade had to include isolates originating from at least two countries, thereby excluding geographically restricted local outbreaks and enriching for lineages with evidence of international dissemination.

### eDCC dilution curve

To assess the extent to which the number of eDCCs among non-DCC strains is influenced by sampling, we performed a simulated analysis of dilution curves for all non-DCC strains. Specifically, we subsampled 50, 100, 300, 500, and 1,000 strains from the total non-DCC strains. For each subset, we calculated the number of the eDCC clades and the proportion of those unclustered strains relative to the overall sampled strains. This process was repeated 20 times at each sampling threshold to ensure the robustness of the results.

### DCCs and eDCCs core genome phylogenies

To minimize the impact of recombination on phylogenetic inference, the assembled whole-genome FASTA files generated using SPAdes^32^ were first annotated using Prokka^43^. The annotated genomes were then subjected to pangenome analysis with Panaroo^44^ to identify orthologous gene clusters across all isolates. Panaroo was run in strict mode to reduce spurious gene annotations and assembly artifacts, and paralog splitting was enabled to improve ortholog discrimination. Core genome alignments were constructed using Panaroo^44^, defining the core genome as genes present in at least 99% of isolates. The resulting core genome alignment was used as input for Gubbins^45^ to detect and remove recombinant regions, thereby generating recombination-free core genome alignments. Core alignments were then obtained using the Gubbins script generate_ska_alignment.py, based on the Panaroo-derived core sequences. Phylogenetic trees for DCCs and eDCCs were reconstructed from recombination-filtered alignments using IQtree2^39^. Maximum-likelihood inference was performed under the GTRCAT model. For computational efficiency, an initial tree was inferred using a simpler substitution model (–first-model JC), followed by refinement under the GTR model (–model GTR) for subsequent iterations.

### Ancestral reconstruction and identification of clade-defining SNPs

For each DCC or eDCC, phylogenetic clusters were first identified based on the topology of the maximum-likelihood tree. To reduce computational burden while preserving genetic diversity, large clusters were pruned using Treemmer^46^, retaining at least 70% of the original tree diversity. These selected isolates were then used for phylogenetic reconstruction and ancestral state inference. We employed SNPParv1.040 ^47^ to infer the sequence of the most recent common ancestor of each DCC/eDCC using general time reversible (GTR) model of nucleotide substitution, with a gamma distribution to account for rate heterogeneity among sites.

Clade-defining SNPs were identified as mutations occurring along the ancestral branch leading to the most recent common ancestor of each DCC/eDCC. Due to the frequent recombination in MAB, we applied a stepwise filtering strategy to improve the specificity of typing SNPs. First, ancestral-branch SNPs shared by multiple DCCs or eDCCs were removed to avoid non-specific clade assignment. Second, candidate SNPs were required to be present in >95% of isolates within the target clade and in < 5% of isolates outside that clade. The resulting clade-defining SNP sets were used as genetic barcodes for DCC/eDCC identification.

### Temporal and spatial transmission dynamics

Temporal phylogenetic reconstructions were performed using the analytical framework described in previous studies^48,49^. Because of the large sample sizes of DCC1–3, the phylogeny was first pruned using Treemmer^46^, removing highly similar strains while preserving approximately 95% of the total tree diversity. We then performed stratified sampling by country, selecting up to approximately 200 isolates from each DCC in each replicate (or all available isolates for DCCs with fewer than 200 isolates). For eDCCs, we focused on clades containing more than 100 isolates and detected in at least three countries, ultimately including eDCC8–14 for further analysis.

For Bayesian Skyline Plot analyses, we used BEAST v.1.10.2^48^ to apply a uniform substitution rate prior ranging from 8.76×10^−8^ – 2.41×10^−7^ substitutions per site per year^16^, employing the HKY nucleotide substitution model with a relaxed log-normal clock and a piecewise constant Bayesian Skyline coalescent prior. Each independent run consisted of 100,000,000 steps, with samples drawn every 10,000 generations. Convergence of Markov Chain Monte Carlo (MCMC) chains and sufficient effective sample sizes (ESS ≥ 200) for all parameters were confirmed using Tracer v.1.7^50^. The resulting skyline plots, visualizing changes in effective population size through time, are presented in Figure 2.

In addition, ancestral geographic states were inferred on the maximum-likelihood phylogeny using PastML^51^. Sampling locations (country level) were assigned as discrete traits for all isolates, and ancestral state reconstruction was performed using the maximum likelihood marginal posterior probability approximation (MPPA) method with an F81-like model. Cross-border transmission events were identified by changes in the inferred geographic state along phylogenetic branches, allowing the reconstruction of international dissemination patterns among lineages.

### Linear regression of time and population size

To test the relationship between time and the population size, we performed linear regression analyses based on the temporal reconstructions inferred using BEAST (v.1.10.2)^48^. For each DCC and eDCC, the effective population sizes and corresponding time estimates were extracted from the Bayesian Skyline Plot (BSP) analyses described above. These BEAST-derived time estimates were then used to evaluate the association between population size and time. Linear regression models were fitted separately for each DCC and eDCC using the linear regression function in R (version 4.2.2).

Additionally, to test whether the potential differences in mutation rates between DCC and Non-DCC strains, as proposed by a recent study^19^, would affect this analysis, we further tested the association using 2.5 SNPs/genome/year for DCCs and 12.0 SNPs/genome/year for eDCCs.

### Mutational events and selective pressure

To investigate genes that have undergone changes in selective pressure within DCCs and eDCCs, we use SNPPar v1.0^47^ to infer the number of independent occurrences of SNPs in the phylogenetic trees of eDCCs and DCCs. SNPPar used the *TreeTime* function for ancestral sequence reconstruction at each node and inferred mutation events along each branch of the phylogenetic tree. We identified all mutations that accumulated prior to the divergence of DCCs or eDCCs as pre-expansion mutations, while those occurring within the DCCs or eDCCs were identified mutations accumulated during expansion. Independent occurrences of the same SNP at the same genomic position across different branches were treated as separate mutational events and retained for subsequent analyses. The SNPs in coding regions were annotated as nonsynonymous or synonymous mutations using SNPPar v1.0^47^. We calculated the pN/pS ratio of mutations from MAB genes using a previous script written in Python^52^. Three separate analyses were then conducted by: (i) combining the mutational events within each subspecies to calculate selection pressure on each gene; and (ii) assessing selective pressure on pre-expansion and during-expansion mutations to identify genes undergoing shifts in selective pressure. For pN/pS analysis, we only inlcuded genes that had more than 15 total mutational events. The genes showing differential selection patterns are reported in Table S4.

### Co-expression cluster identification and mutation profiling

The transcriptomic expression matrix (RPKM) was obtained from the dataset curated by Bei et al^20^. During preprocessing, only genes with a cross-sample variance greater than 1 were retained. The filtered matrix was subjected to quantile normalization^53^ (using the qnorm package, v.0.9.0), followed by a log2 transformation with a pseudo-count of 1 [*log2(normalized RPKM + 1)*]. Subsequently, batch effects were corrected using the conorm package (v.1.2.0), with the original study accession treated as the batch covariate. The batch-corrected matrix was utilized to calculate pairwise Pearson correlation coefficients for 55 positively selected genes to construct a correlation matrix. Hierarchical clustering was then performed on this matrix based on cosine distance using the complete linkage method. The dendrogram was cut at a threshold corresponding to 70% of the maximum clustering distance to define co-expression modules.

Baseline annotation information was first obtained using the Mycobrowser database^54^ (Release 5, 2024-07-11) with MAB ATCC19977 as reference strain. For genes with incomplete annotations or unknown functions, their amino acid sequences were retrieved from the UniProt database^55^ (Taxon ID: 561007). These sequences were initially subjected to orthology-based functional prediction using the eggNOG-mapper web server^56^ (http://eggnog-mapper.embl.de). Subsequently, structural searches were conducted using the Foldseek web server^57^ against the AlphaFold Protein Structure Database (AFDB)^58^. We prioritized recording high-ranking structural matches with *Mycobacterium tuberculosis* H37Rv proteins; if unavailable, high-ranking matches from *Mycobacterium smegmatis* were recorded, and if still unavailable, *Escherichia coli* matches were selected. These structural similarities served as auxiliary evidence to aid in inferring potential homologous relationships and functional clues. Furthermore, we conducted a manual review of relevant publications retrieved from the PubMed database, focusing on these 55 positively selected genes and reports of their homologs in *Mycobacterium tuberculosis* and *Mycobacterium smegmatis*. Based on the integrated sequence, structural, and literature evidence, the target genes were manually assigned to the following primary functional categories: transcriptional regulation, cell envelope, cAMP metabolism, ribosome function, redox-related, DNA replication, lipid metabolism, energy metabolism, and macrolide resistance. During visualization, functional genes associated with environmental regulation and cAMP metabolism were specifically highlighted. The functional consequences of non-synonymous SNPs were annotated, and the relative frequencies of loss-of-function (LOF) variants were calculated. The LOF relative frequency for an individual gene was defined as the number of non-synonymous SNPs annotated as LOF within that gene, divided by the total number of annotated non-synonymous SNPs within the same gene. All data analyses and visualizations were performed using Python (v.3.11.14) with the seaborn (v.0.13.2) and matplotlib (v.3.10.8) packages.

### Data Availability

Raw data and all analyzing scripts in the article have been uploaded to GitHub (https://github.com/zhuchendi0520/MABC_eDCCs.git). The raw data include: 1) FASTA files containing aligned and concatenated DNA sequences used for constructing the phylogenetic trees in Figures 1C–F, 2B, and 5C; 2) Gubbins tree files for each ECC and DCC; and 3) XML files for all BEAST2 temporal and geographic reconstructions.

## Acknowledgment

We thank Howard Takiff for assistance in editing the manuscript.

## Reference

1. Moore, M. & Frerichs, J.B. An unusual acid-fast infection of the knee with subcutaneous, abscess-like lesions of the gluteal region; report of a case with a study of the organism, Mycobacterium abscessus, n. sp. J Invest Dermatol 20, 133–169 (1953).

2. Tortoli, E., et al. Emended description of Mycobacterium abscessus, Mycobacterium abscessus subsp. abscessus and Mycobacteriumabscessus subsp. bolletii and designation of Mycobacteriumabscessus subsp. massiliense comb. nov. Int J Syst Evol Microbiol 66, 4471–4479 (2016).

3. Kwak, N., et al. M ycobacterium abscessus pulmonary disease: individual patient data meta-analysis. Eur Respir J 54(2019).

4. van Ingen, J., Boeree, M.J., van Soolingen, D. & Mouton, J.W. Resistance mechanisms and drug susceptibility testing of nontuberculous mycobacteria. Drug Resist Updat 15, 149–161 (2012).

5. Koh, W.J., et al. Mycobacterial Characteristics and Treatment Outcomes in Mycobacterium abscessus Lung Disease. Clin Infect Dis 64, 309–316 (2017).

6. Falkinham, J.O., 3rd. Environmental sources of nontuberculous mycobacteria. Clin Chest Med 36, 35–41 (2015).

7. Thomson, R., et al. Isolation of nontuberculous mycobacteria (NTM) from household water and shower aerosols in patients with pulmonary disease caused by NTM. J Clin Microbiol 51, 3006–3011 (2013).

8. Bryant, J.M., et al. Whole-genome sequencing to identify transmission of Mycobacterium abscessus between patients with cystic fibrosis: a retrospective cohort study. Lancet 381, 1551–1560 (2013).

9. Gross, J.E., et al. Investigating Nontuberculous Mycobacteria Transmission at the Colorado Adult Cystic Fibrosis Program. Am J Respir Crit Care Med 205, 1064–1074 (2022).

10. Lipworth, S., et al. Epidemiology of Mycobacterium abscessus in England: an observational study. Lancet Microbe 2, e498–e507 (2021).

11. Bryant, J.M., et al. Stepwise pathogenic evolution of Mycobacterium abscessus. Science 372(2021).

12. Bryant, J.M., et al. Emergence and spread of a human-transmissible multidrug-resistant nontuberculous mycobacterium. Science 354, 751–757 (2016).

13. Thomson, R.M., et al. Infection by Clonally Related Mycobacterium abscessus Isolates: The Role of Drinking Water. Am J Respir Crit Care Med 211, 842–853 (2025).

14. Li, X., et al. Population genetic analysis of clinical Mycobacterium abscessus complex strains in China. Front Cell Infect Microbiol 14, 1496896 (2024).

15. Yoshida, M., et al. Molecular Epidemiological Characteristics of Mycobacterium abscessus Complex Derived from Non-Cystic Fibrosis Patients in Japan and Taiwan. Microbiol Spectr 10, e0057122 (2022).

16. Ruis, C., et al. Dissemination of Mycobacterium abscessus via global transmission networks. Nat Microbiol 6, 1279–1288 (2021).

17. Johansen, M.D., Herrmann, J.L. & Kremer, L. Non-tuberculous mycobacteria and the rise of Mycobacterium abscessus. Nat Rev Microbiol 18, 392–407 (2020).

18. Victoria, L., Gupta, A., Gómez, J.L. & Robledo, J. Mycobacterium abscessus complex: A Review of Recent Developments in an Emerging Pathogen. Front Cell Infect Microbiol 11, 659997 (2021).

19. Commins, N., et al. Mutation rates and adaptive variation among the clinically dominant clusters of Mycobacterium abscessus. Proc Natl Acad Sci U S A 120, e2302033120 (2023).

20. Bei, C., et al. Genetically encoded transcriptional plasticity underlies stress adaptation in Mycobacterium tuberculosis. Nat Commun 15, 3088 (2024).

21. Everall, I., et al. Genomic epidemiology of a national outbreak of post-surgical Mycobacterium abscessus wound infections in Brazil. Microb Genom 3, e000111 (2017).

22. Planet, P.J., et al. Architecture of a Species: Phylogenomics of Staphylococcus aureus. Trends Microbiol 25, 153–166 (2017).

23. Weimann, A., et al. Evolution and host-specific adaptation of Pseudomonas aeruginosa. Science 385, eadi0908 (2024).

24. Yu, X.A., et al. Genome-wide sweeps create ecological units in the human gut microbiome. Nature 655, 202–209 (2026).

25. Catherinot, E., et al. Mycobacterium avium and Mycobacterium abscessus complex target distinct cystic fibrosis patient subpopulations. J Cyst Fibros 12, 74–80 (2013).

26. Turenne, C.Y. Nontuberculous mycobacteria: Insights on taxonomy and evolution. Infect Genet Evol 72, 159–168 (2019).

27. Tortoli, E., et al. The new phylogeny of the genus Mycobacterium: The old and the news. Infect Genet Evol 56, 19–25 (2017).

28. Poerio, N., et al. Combined Host- and Pathogen-Directed Therapy for the Control of Mycobacterium abscessus Infection. Microbiol Spectr 10, e0254621 (2022).

29. Touré, H., et al. Mycobacterium abscessus Opsonization Allows an Escape from the Defensin Bactericidal Action in Drosophila. Microbiol Spectr 11, e0077723 (2023).

30. Winters, C.G., et al. Disulfiram Is Effective against Drug-Resistant Mycobacterium abscessus in a Zebrafish Embryo Infection Model. Antimicrob Agents Chemother 66, e0053922 (2022).

31. Lipworth, S., et al. Whole-Genome Sequencing for Predicting Clarithromycin Resistance in Mycobacterium abscessus. Antimicrob Agents Chemother 63(2019).

32. Bankevich, A., et al. SPAdes: a new genome assembly algorithm and its applications to single-cell sequencing. J Comput Biol 19, 455–477 (2012).

33. Hernández-Salmerón, J.E. & Moreno-Hagelsieb, G. FastANI, Mash and Dashing equally differentiate between Klebsiella species. PeerJ 10, e13784 (2022).

34. Joshi NA, Fass JN. (2011). Sickle: A sliding-window, adaptive, quality-based trimming tool for FastQ files (Version 1.33) [Software]. Available at https://github.com/najoshi/sickle.

35. Jung, Y. & Han, D. BWA-MEME: BWA-MEM emulated with a machine learning approach. Bioinformatics 38, 2404–2413 (2022).

36. Li, H., et al. The Sequence Alignment/Map format and SAMtools. Bioinformatics 25, 2078–2079 (2009).

37. Koboldt, D.C., et al. VarScan 2: somatic mutation and copy number alteration discovery in cancer by exome sequencing. Genome Res 22, 568–576 (2012).

38. Price, M.N., Dehal, P.S. & Arkin, A.P. FastTree: computing large minimum evolution trees with profiles instead of a distance matrix. Mol Biol Evol 26, 1641–1650 (2009).

39. Minh, B.Q., et al. IQ-TREE 2: New Models and Efficient Methods for Phylogenetic Inference in the Genomic Era. Mol Biol Evol 37, 1530–1534 (2020).

40. Kalyaanamoorthy, S., Minh, B.Q., Wong, T.K.F., von Haeseler, A. & Jermiin, L.S. ModelFinder: fast model selection for accurate phylogenetic estimates. Nat Methods 14, 587–589 (2017).

41. Rambaut, A. FigTree v1.4.4. (Institute of Evolutionary Biology, University of Edinburgh, 2018).

42. Letunic, I. & Bork, P. Interactive tree of life (iTOL) v3: an online tool for the display and annotation of phylogenetic and other trees. Nucleic Acids Res 44, W242–245 (2016).

43. Seemann, T. Prokka: rapid prokaryotic genome annotation. Bioinformatics 30, 2068–2069 (2014).

44. Tonkin-Hill, G., et al. Producing polished prokaryotic pangenomes with the Panaroo pipeline. Genome Biol 21, 180 (2020).

45. Croucher, N.J., et al. Rapid phylogenetic analysis of large samples of recombinant bacterial whole genome sequences using Gubbins. Nucleic Acids Res 43, e15 (2015).

46. Menardo, F., et al. Treemmer: a tool to reduce large phylogenetic datasets with minimal loss of diversity. Bmc Bioinformatics 19, 164 (2018).

47. Edwards, D.J., Duchene, S., Pope, B. & Holt, K.E. SNPPar: identifying convergent evolution and other homoplasies from microbial whole-genome alignments. Microb Genom 7(2021).

48. Bouckaert, R., et al. BEAST 2: a software platform for Bayesian evolutionary analysis. PLoS Comput Biol 10, e1003537 (2014).

49. Menardo, F., Duchene, S., Brites, D. & Gagneux, S. The molecular clock of Mycobacterium tuberculosis. PLoS pathogens 15, e1008067 (2019).

50. Rambaut, A., Drummond, A.J., Xie, D., Baele, G. & Suchard, M.A. Posterior Summarization in Bayesian Phylogenetics Using Tracer 1.7. Syst Biol 67, 901–904 (2018).

51. Ishikawa, S.A., Zhukova, A., Iwasaki, W. & Gascuel, O. A Fast Likelihood Method to Reconstruct and Visualize Ancestral Scenarios. Mol Biol Evol 36, 2069–2085 (2019).

52. Trauner, A., et al. The within-host population dynamics of Mycobacterium tuberculosis vary with treatment efficacy. Genome Biol 18, 71 (2017).

53. Bolstad, B.M., Irizarry, R.A., Astrand, M. & Speed, T.P. A comparison of normalization methods for high density oligonucleotide array data based on variance and bias. Bioinformatics 19, 185–193 (2003).

54. Kapopoulou, A., Lew, J.M. & Cole, S.T. The MycoBrowser portal: a comprehensive and manually annotated resource for mycobacterial genomes. Tuberculosis (Edinb) 91, 8–13 (2011).

55. UniProt: the Universal Protein Knowledgebase in 2025. Nucleic Acids Res 53, D609–d617 (2025).

56. Cantalapiedra, C.P., Hernández-Plaza, A., Letunic, I., Bork, P. & Huerta-Cepas, J. eggNOG-mapper v2: Functional Annotation, Orthology Assignments, and Domain Prediction at the Metagenomic Scale. Mol Biol Evol 38, 5825–5829 (2021).

57. van Kempen, M., et al. Fast and accurate protein structure search with Foldseek. Nat Biotechnol 42, 243–246 (2024).

58. Varadi, M., et al. AlphaFold Protein Structure Database in 2024: providing structure coverage for over 214 million protein sequences. Nucleic Acids Res 52, D368–d375 (2024).

