## Supplementary Figures for "Parallel Emergence and Adaptive Evolution of Multicountry Circulating *Mycobacterium abscessus* Clones"

Chendi Zhu, et al.

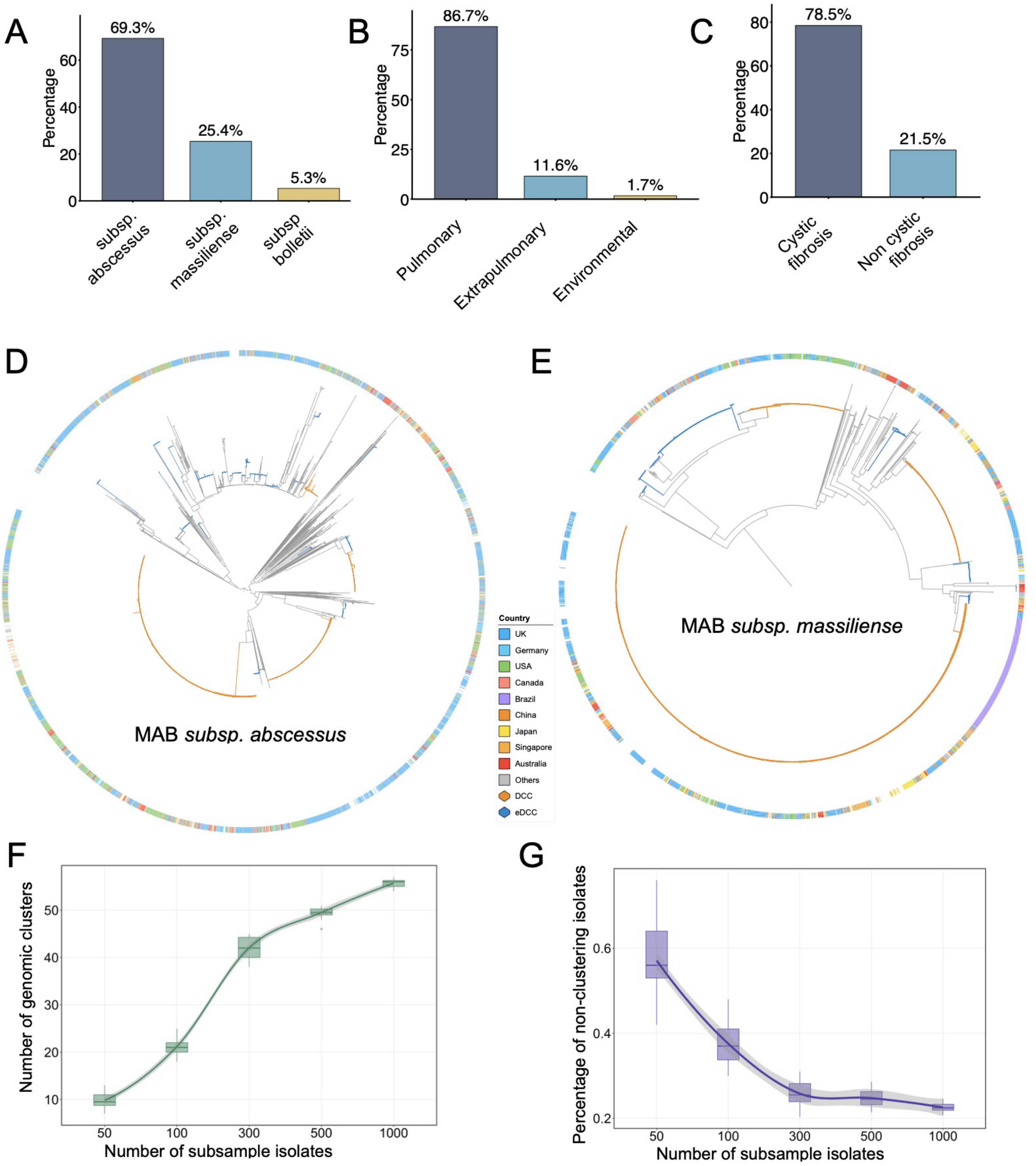


**Supplementary Figure 1. Global genomic composition and sampling characteristics of the MAB population.** (**A**) Relative abundance of the three MAB subspecies in the global genomic dataset, as determined by average nucleotide identity (ANI). (**B**) Distribution of isolates by source, with most isolates derived from pulmonary specimens. (**C**) Distribution of clinical isolates by host cystic fibrosis (CF) status. (**D–E**) Circular phylogenies of MAB subsp. *abscessus* (**D**) and subsp. *massiliense* (**E**). The outer ring indicates the country of isolation, while orange and blue branches denote DCCs and eDCCs, respectively. (**F**) Subsampling analysis showing the relationship between the number of isolates included and the number of genomic clusters identified. (**G**) Relationship between subsample size and the proportion of isolates not assigned to any genomic cluster.

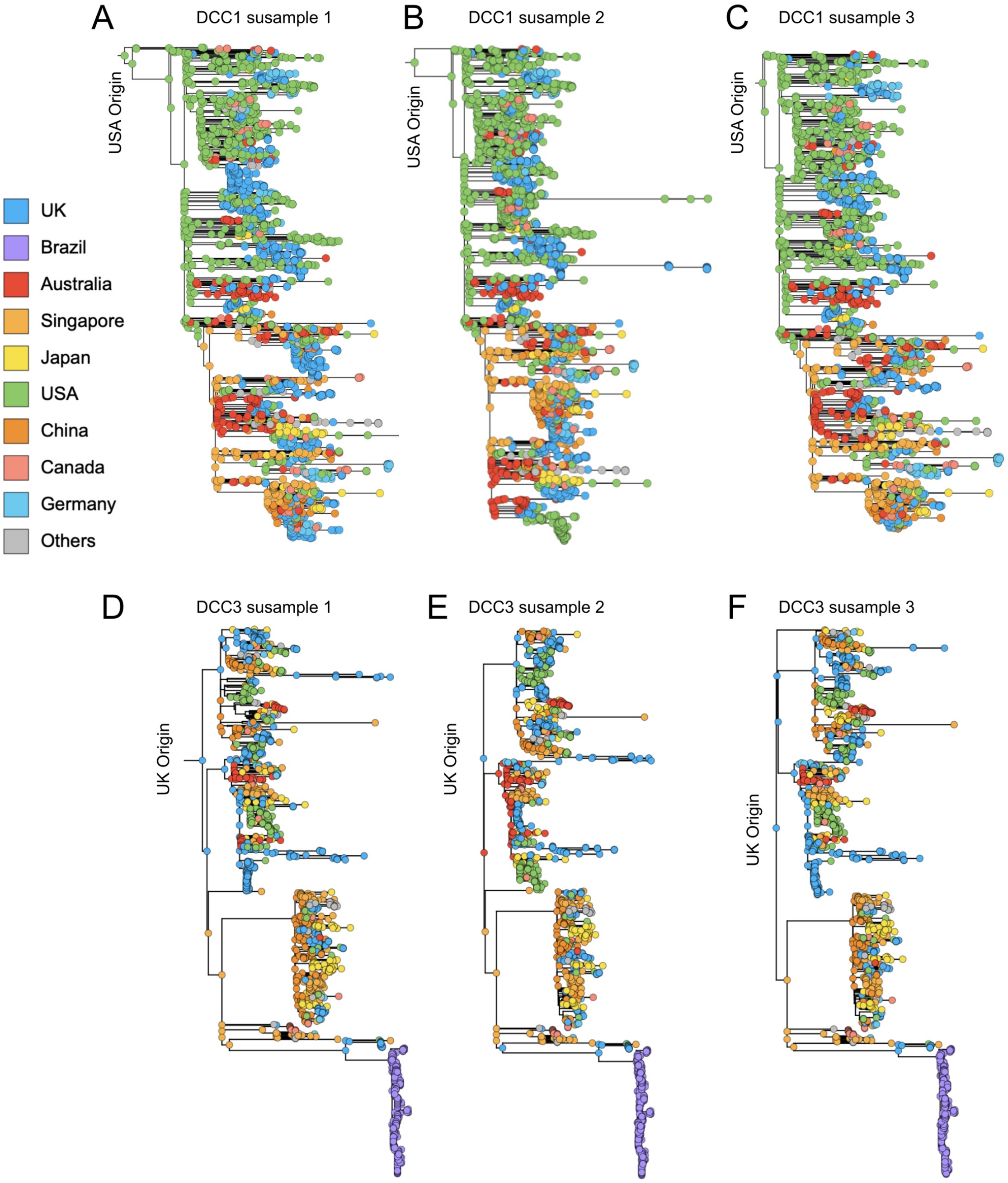


**Supplementary Figure 2. Phylogeographic reconstruction using subsampled datasets with balanced country representation.** (**A–C**) Phylogeographic reconstruction of the DCC1 clade based on three independently generated subsampled datasets with balanced country representation to control for potential sampling bias. (**D–F**) Phylogeographic reconstruction of the DCC3 clade based on the same subsampling strategy.

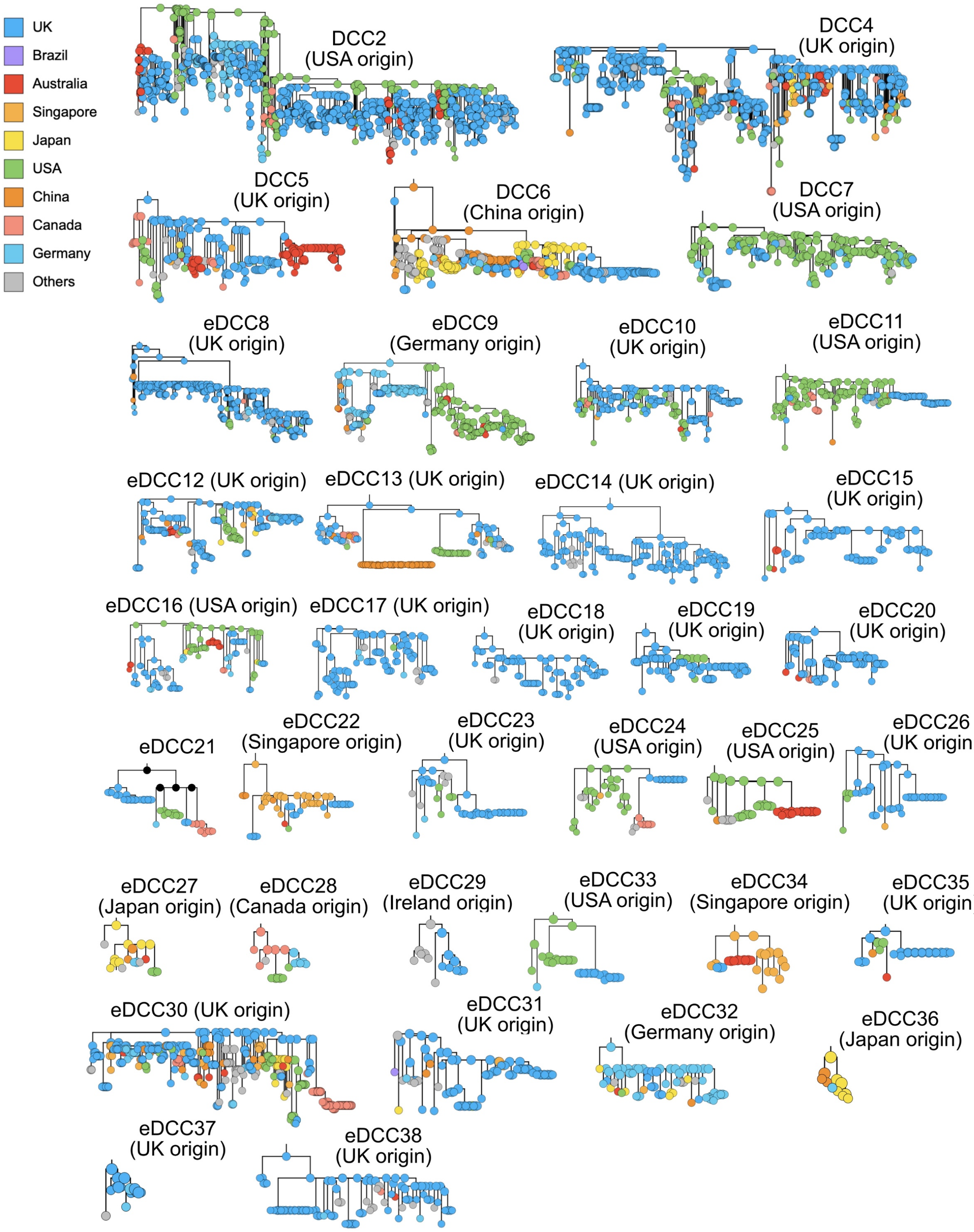


**Supplementary Figure 3. Country distribution of isolates across DCC and eDCC phylogenies.** Individual phylogenies are shown for established DCCs and newly identified eDCCs. Tips are colored according to the country of sampling, and the inferred geographic origin of each clade is indicated in parentheses when available. The trees show the geographic composition and fine-scale phylogenetic structure of each circulating clone, including both country-restricted expansions and internationally distributed lineages.

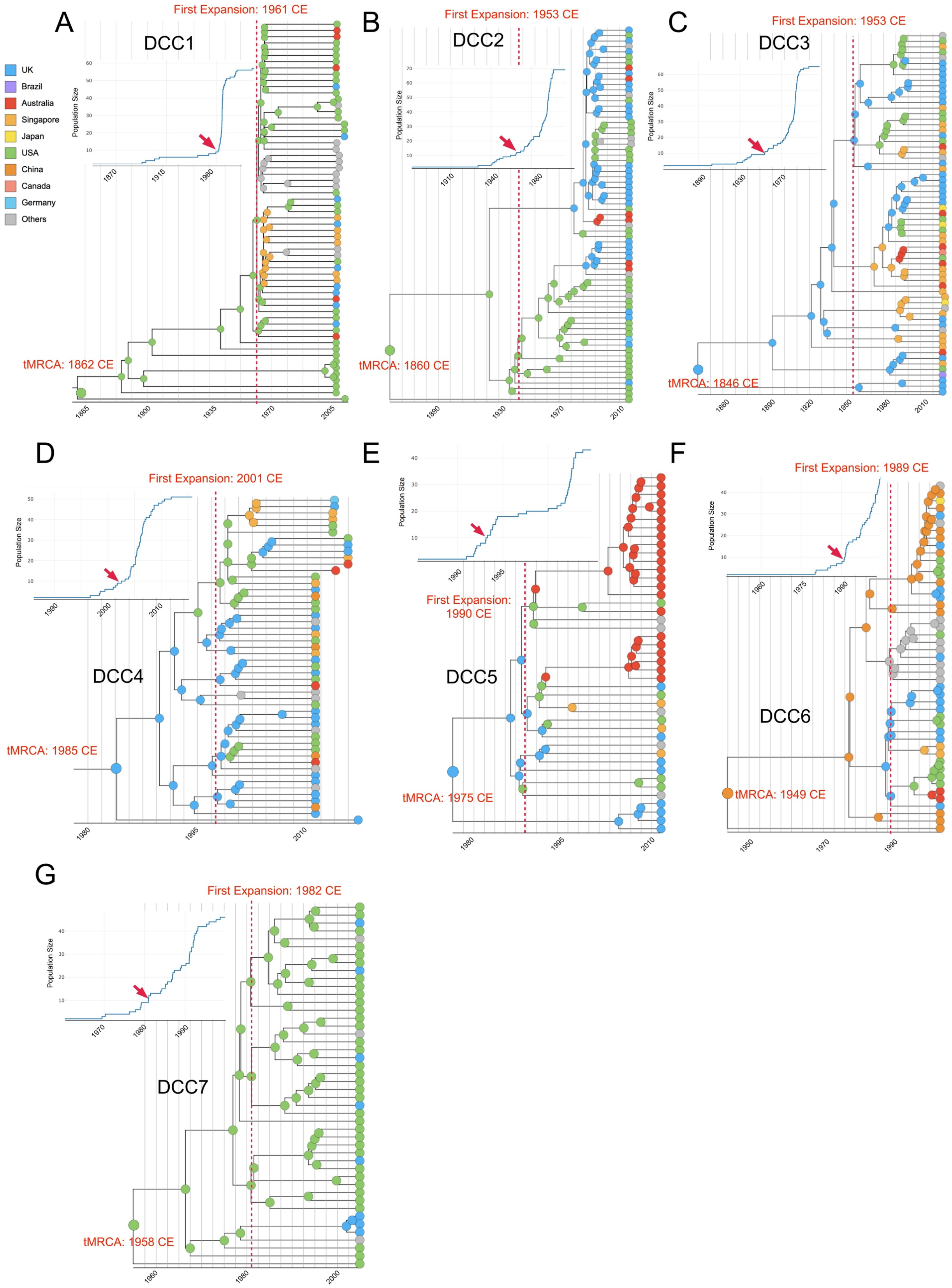


**Supplementary Figure 4. Time-calibrated phylogenies of major DCC and eDCC clades inferred using BEAST.** (**A–G**) Time-calibrated phylogenetic trees of the seven previously described dominant circulating clones (DCC1–DCC7). Insets show the cumulative number of phylogenetic nodes over time, with arrows indicating the onset of rapid clade expansion. Dashed red lines indicate the estimated timing of the first expansion, and the estimated time to the most recent common ancestor (tMRCA) is shown for each clade. (**H–N**) Time-calibrated phylogenetic trees of eDCC8–eDCC14, displayed using the same format.

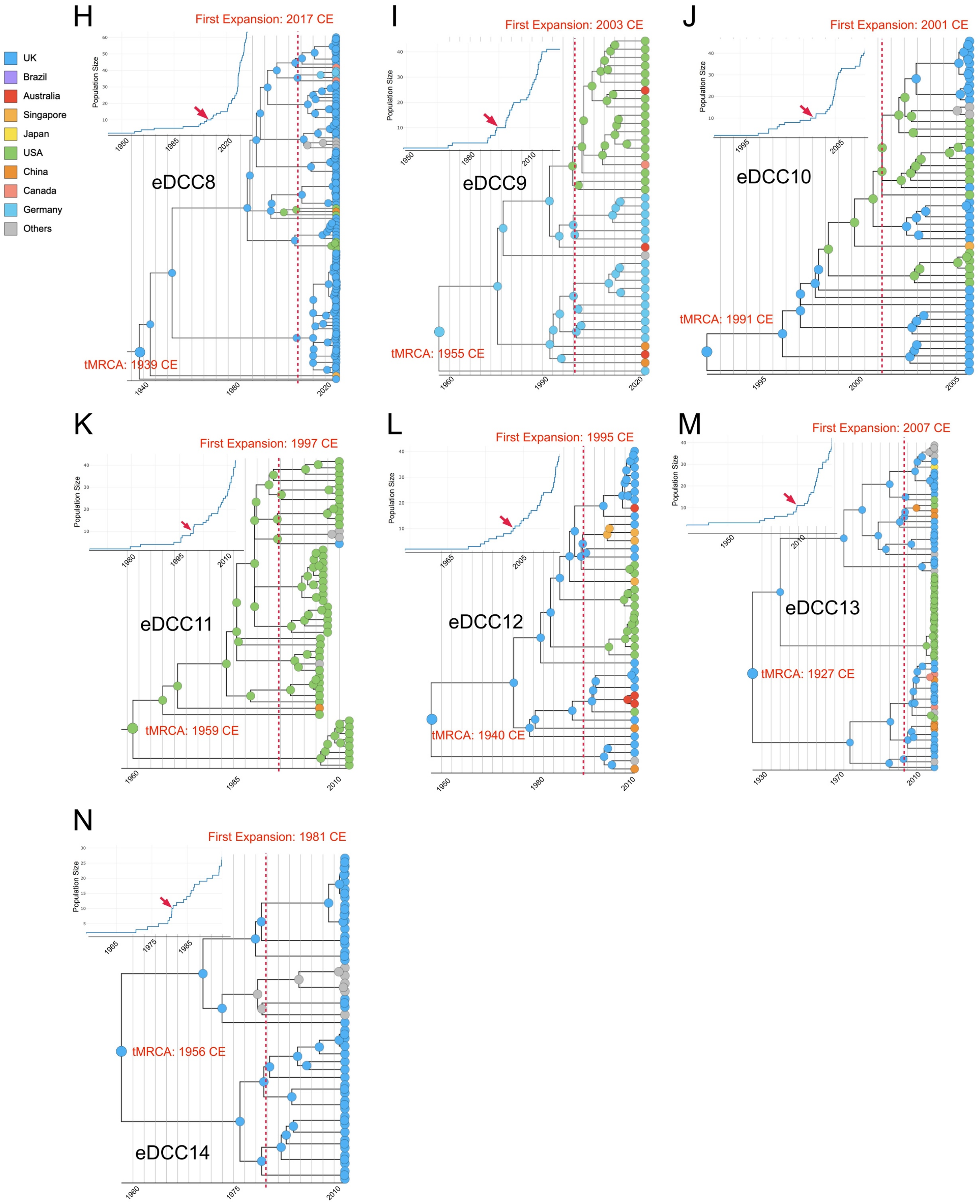


**Supplementary Figure 4 (continued). Time-calibrated phylogenies of major DCC and eDCC clades inferred using BEAST.** (**A–G**) Time-calibrated phylogenetic trees of the seven previously described dominant circulating clones (DCC1–DCC7). Insets show the cumulative number of phylogenetic nodes over time, with arrows indicating the onset of rapid clade expansion. Dashed red lines indicate the estimated timing of the first expansion, and the estimated time to the most recent common ancestor (tMRCA) is shown for each clade. (**H–N**) Time-calibrated phylogenetic trees of eDCC8–eDCC14, displayed using the same format.

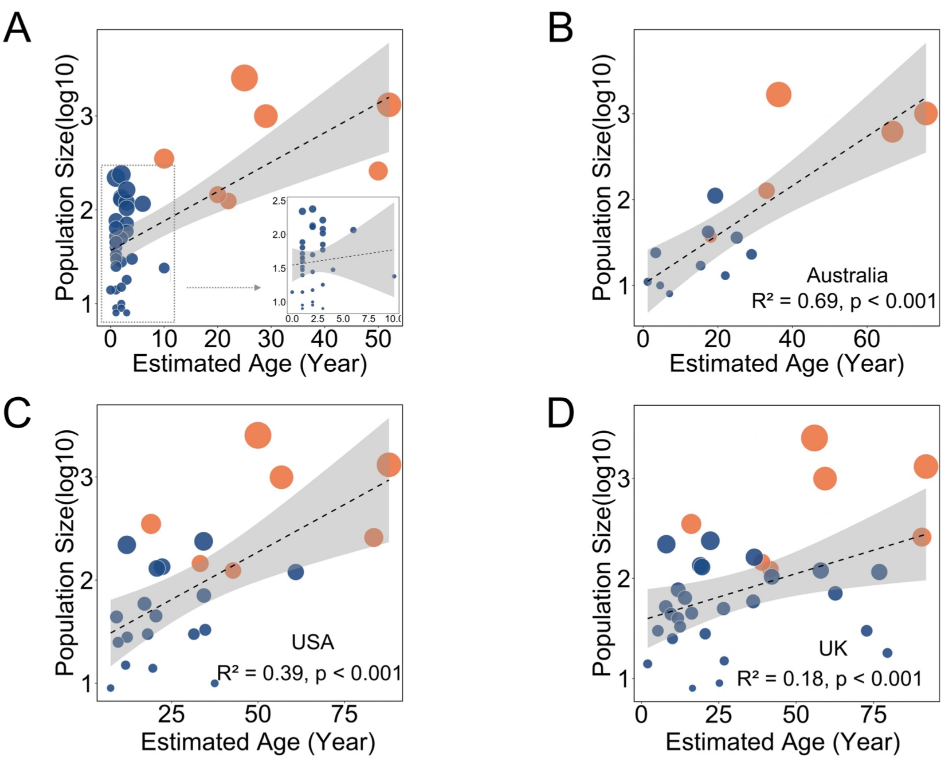


**Supplementary Figure 5. Correlation between clade age and population size**. (**A**) Correlation between estimated clade age and population size using a recently proposed clade-specific mutation rate. Each point represents one clade, with point size proportional to population size. The dashed line indicates the fitted linear regression, and the shaded area represents the 95% confidence interval ($R^2 = 0.43$, $P < 0.001$). The inset shows an enlarged view of younger clades. (**B–D**) Corresponding analyses restricted to isolates sampled from Australia (**B**), the USA (**C**), and the UK (**D**).

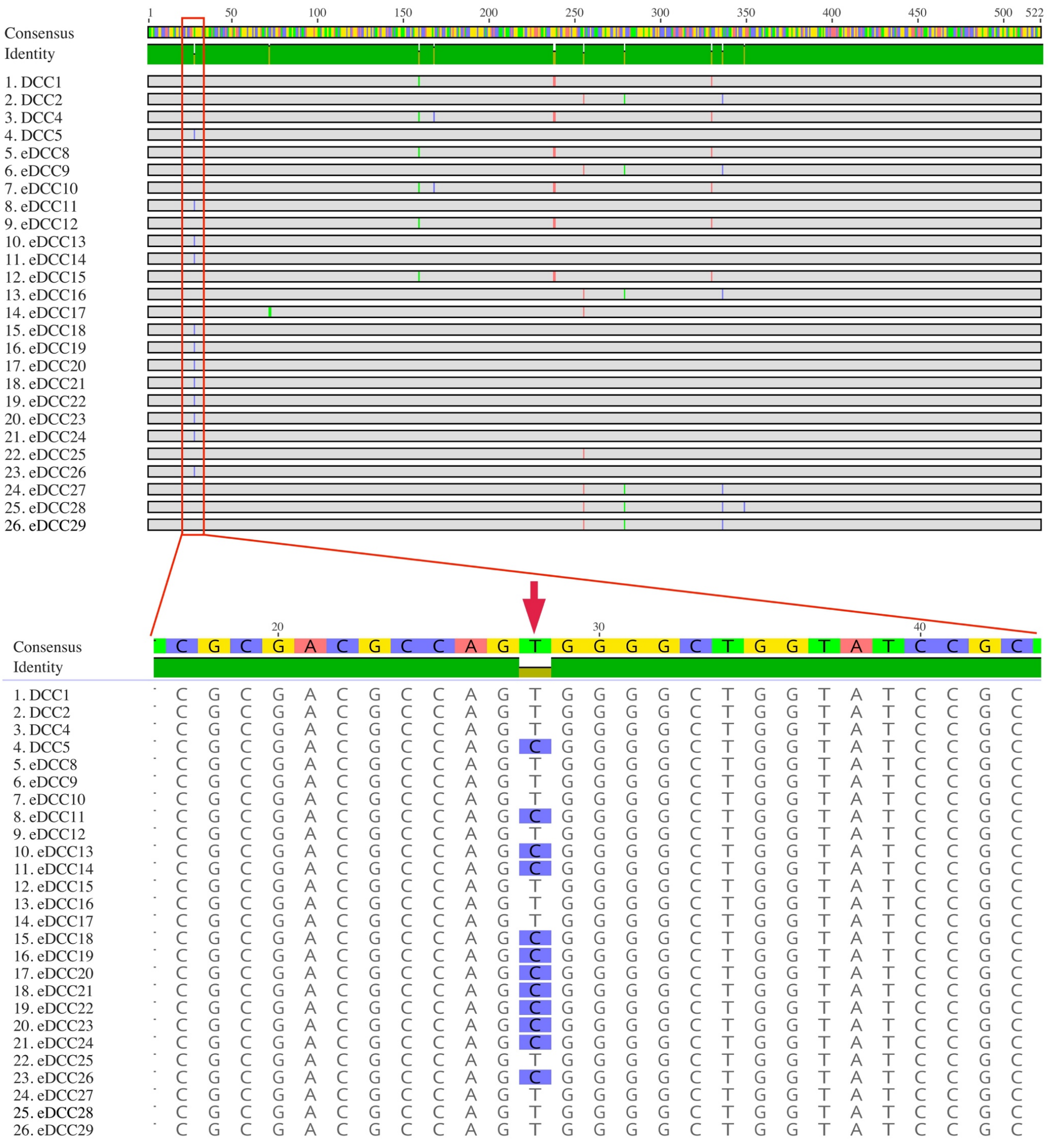


**Supplementary Figure 6. Sequence alignment of *erm(41)* from representative MAB subsp*. abscessus* circulating clades**. The lower panel highlights nucleotide position 28, the canonical T28/C28 polymorphism associated with inducible macrolide resistance, showing its distribution across representative DCC and eDCC lineages.

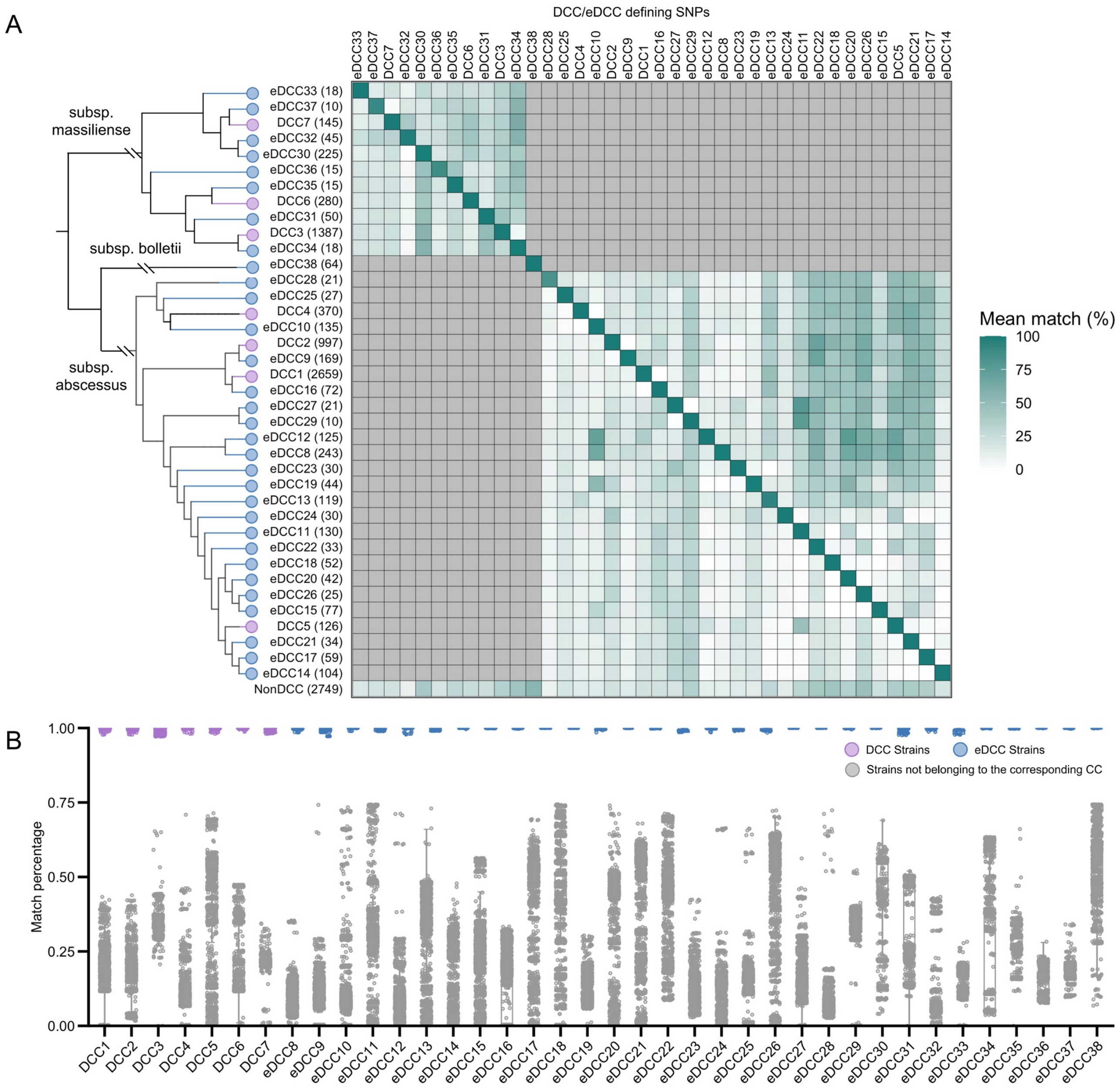


**Supplementary Figure 7. Validation and optimization of DCC- and eDCC-defining barcode SNPs**. (**A**) Cross-clade matching of clade-defining barcode SNPs. The heatmap shows the mean percentage of barcode SNPs matched by isolates from each DCC, eDCC, or non-DCC group, with gray shading indicating comparisons between different subspecies. The phylogeny on the left shows the relationships among the evaluated clades, and numbers in parentheses indicate the number of isolates in each group. (**B**) Distribution of barcode-matching percentages for individual isolates against each clade-specific barcode set. Isolates belonging to the corresponding DCC or eDCC are shown in purple or blue, respectively, whereas isolates not belonging to the corresponding clade are shown in gray. (**C**) Genomic distribution of shared barcode SNPs, highlighting loci shared between clades that were excluded from the final barcode sets because they are potentially attributable to historical recombination (see following pages).

C

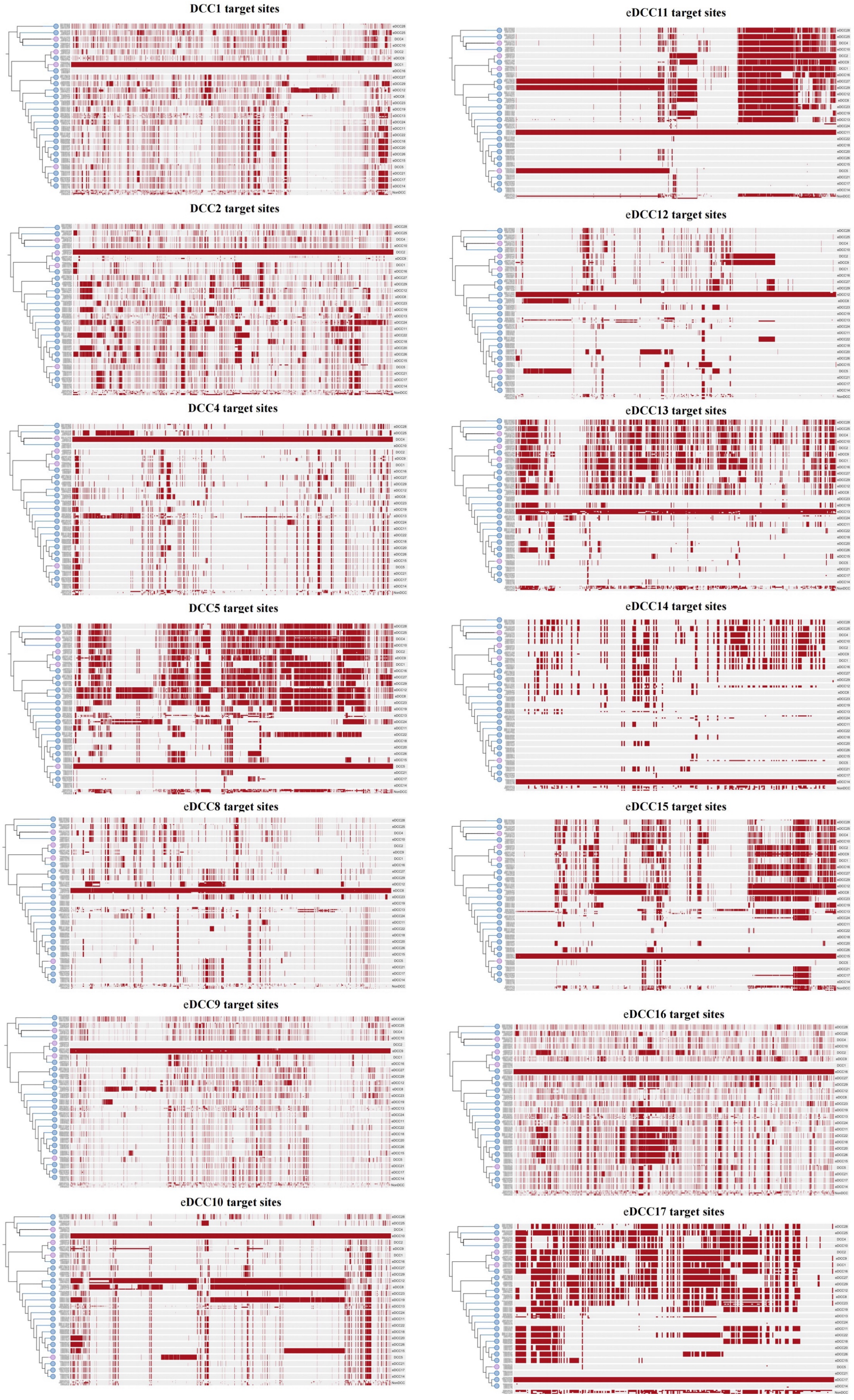
DCC1 target sites
eDCC11 target sites
DCC2 target sites
eDCC12 target sites
DCC4 target sites
eDCC13 target sites
DCC5 target sites
eDCC14 target sites
eDCC8 target sites
eDCC15 target sites
eDCC9 target sites
eDCC16 target sites
eDCC10 target sites
eDCC17 target sites

eDCC18 target sites

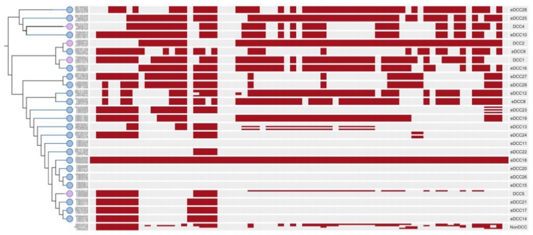

eDCC19target sites

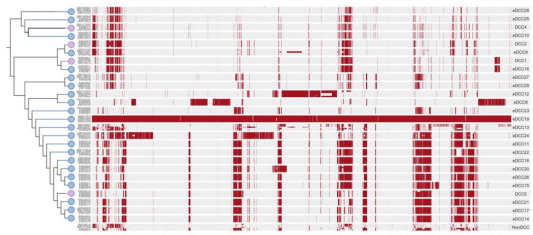

eDCC20 target sites

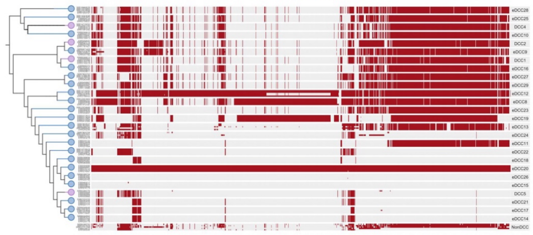

eDCC21 target sites

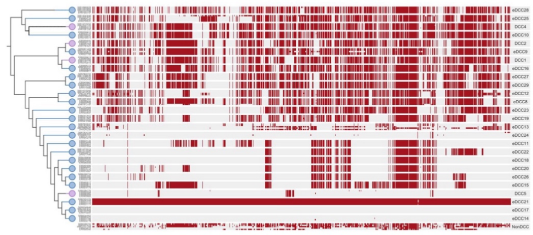

eDCC22 target sites

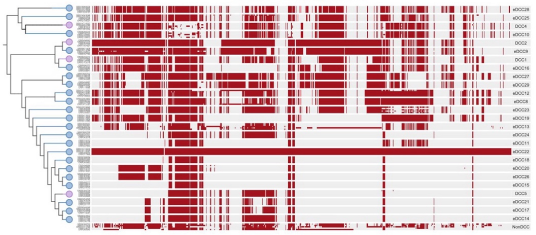

eDCC23 target sites

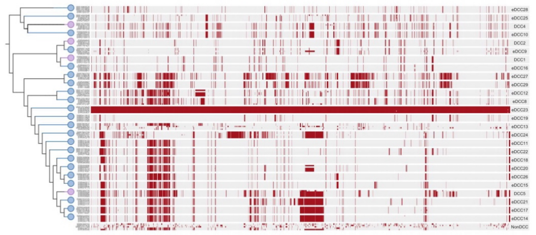

eDCC24 target sites

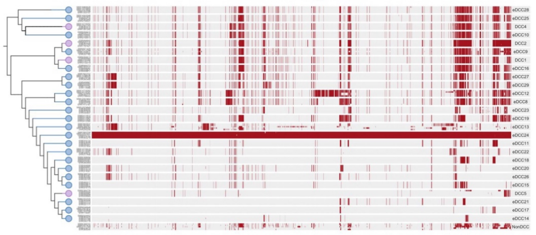

eDCC25 target sites

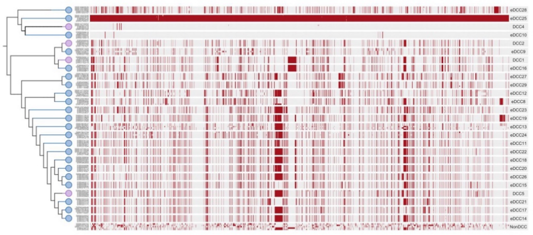

eDCC26 target sites

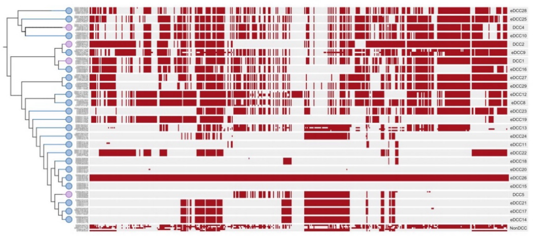

eDCC27 target sites

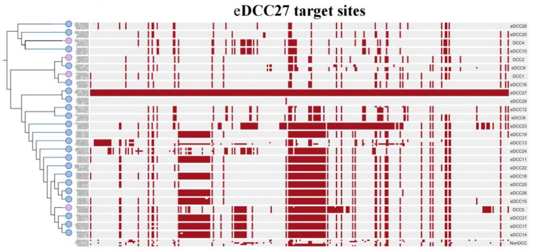

eDCC28 target sites

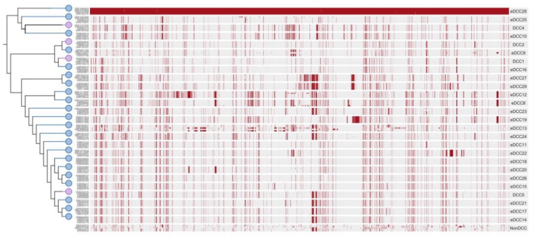

eDCC29target sites

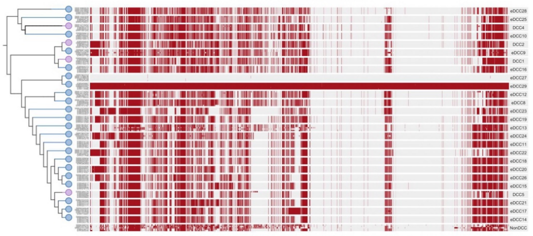

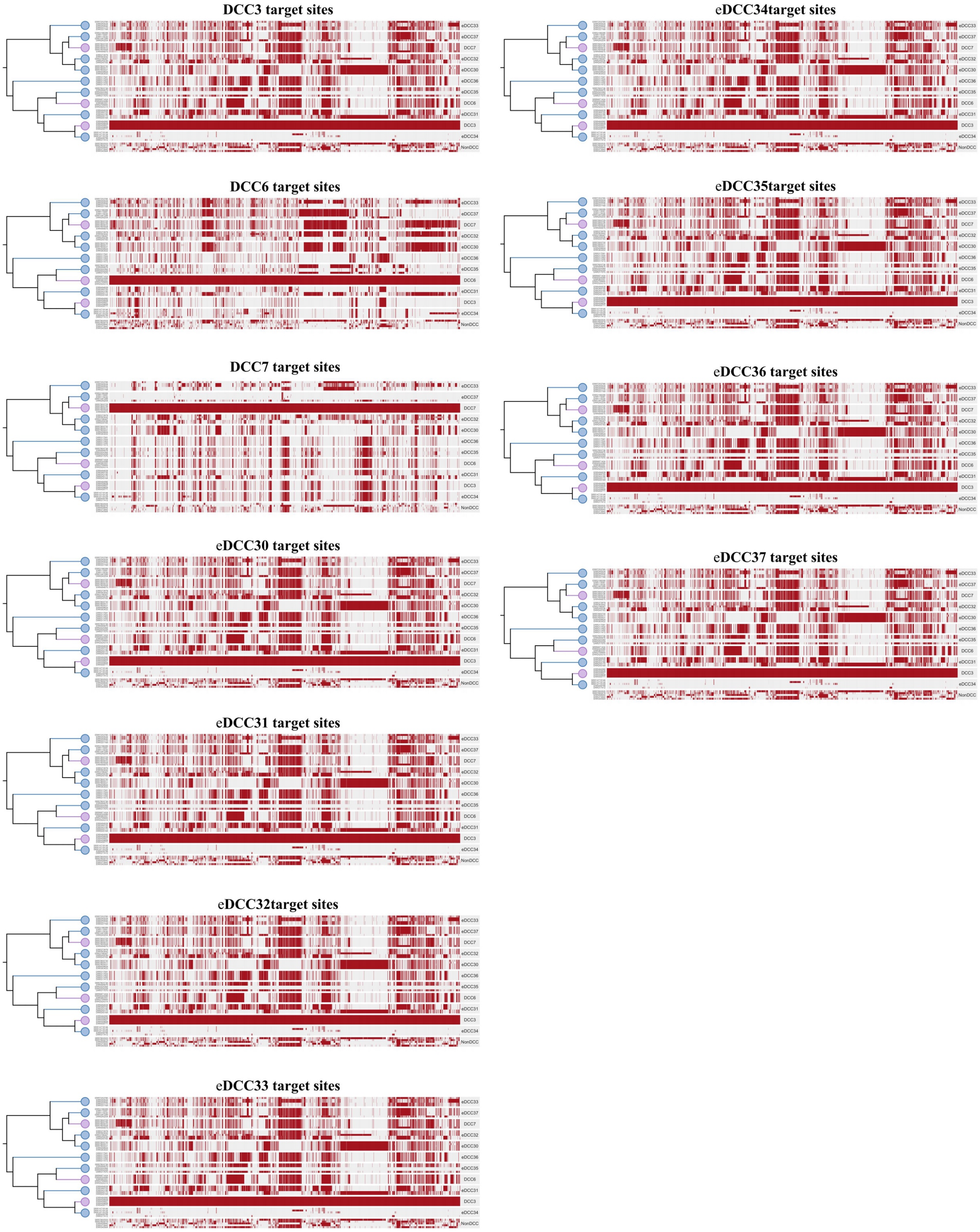


**Supplementary Figure 7 (continued). Validation and optimization of DCC- and eDCC-defining barcode SNPs**. (**A**) Cross-clade matching of clade-defining barcode SNPs. The heatmap shows the mean percentage of barcode SNPs matched by isolates from each DCC, eDCC, or non-DCC group, with gray shading indicating comparisons between different subspecies. The phylogeny on the left shows the relationships among the evaluated clades, and numbers in parentheses indicate the number of isolates in each group. (**B**) Distribution of barcode-matching percentages for individual isolates against each clade-specific barcode set. Isolates belonging to the corresponding DCC or eDCC are shown in purple or blue, respectively, whereas isolates not belonging to the corresponding clade are shown in gray. (**C**) Genomic distribution of shared barcode SNPs, highlighting loci shared between clades that were excluded from the final barcode sets because they are potentially attributable to historical recombination.

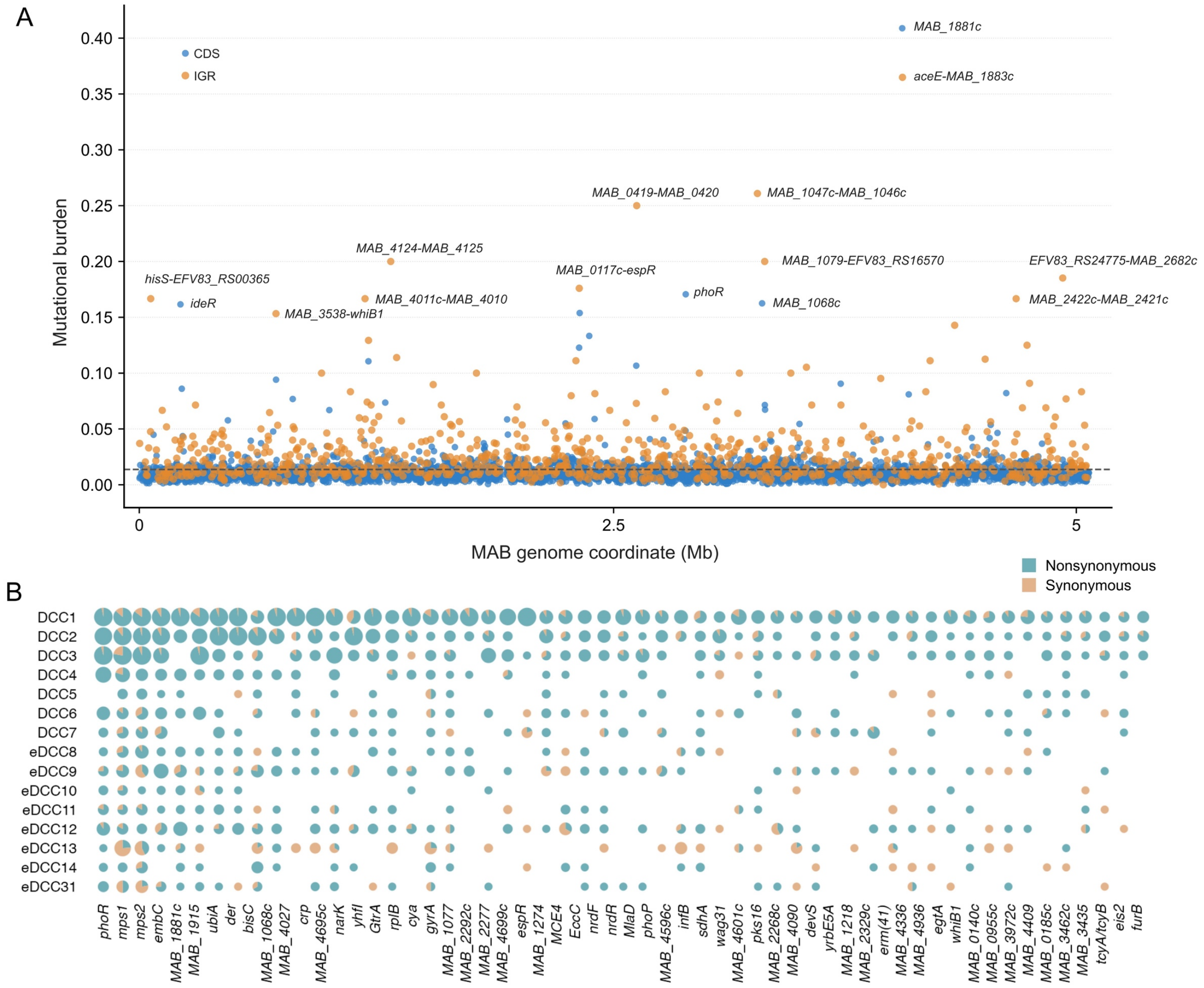


**Supplementary Figure 8. Mutation profiles of positively selected genes across DCCs and eDCCs**. (**A**) Genome-wide distribution of mutational burden across the MAB genome (GZ002). Blue and orange points represent coding sequences (CDSs) and intergenic regions (IGRs), respectively. The dashed line indicates the genome-wide average mutational burden. (**B**) Bubble plot showing the distribution of synonymous and nonsynonymous mutations in positively selected genes across representative DCCs and eDCCs. Circle size reflects the number of mutations observed for each gene–clade combination, and color indicates mutation type, with teal representing nonsynonymous mutations and orange representing synonymous mutations.

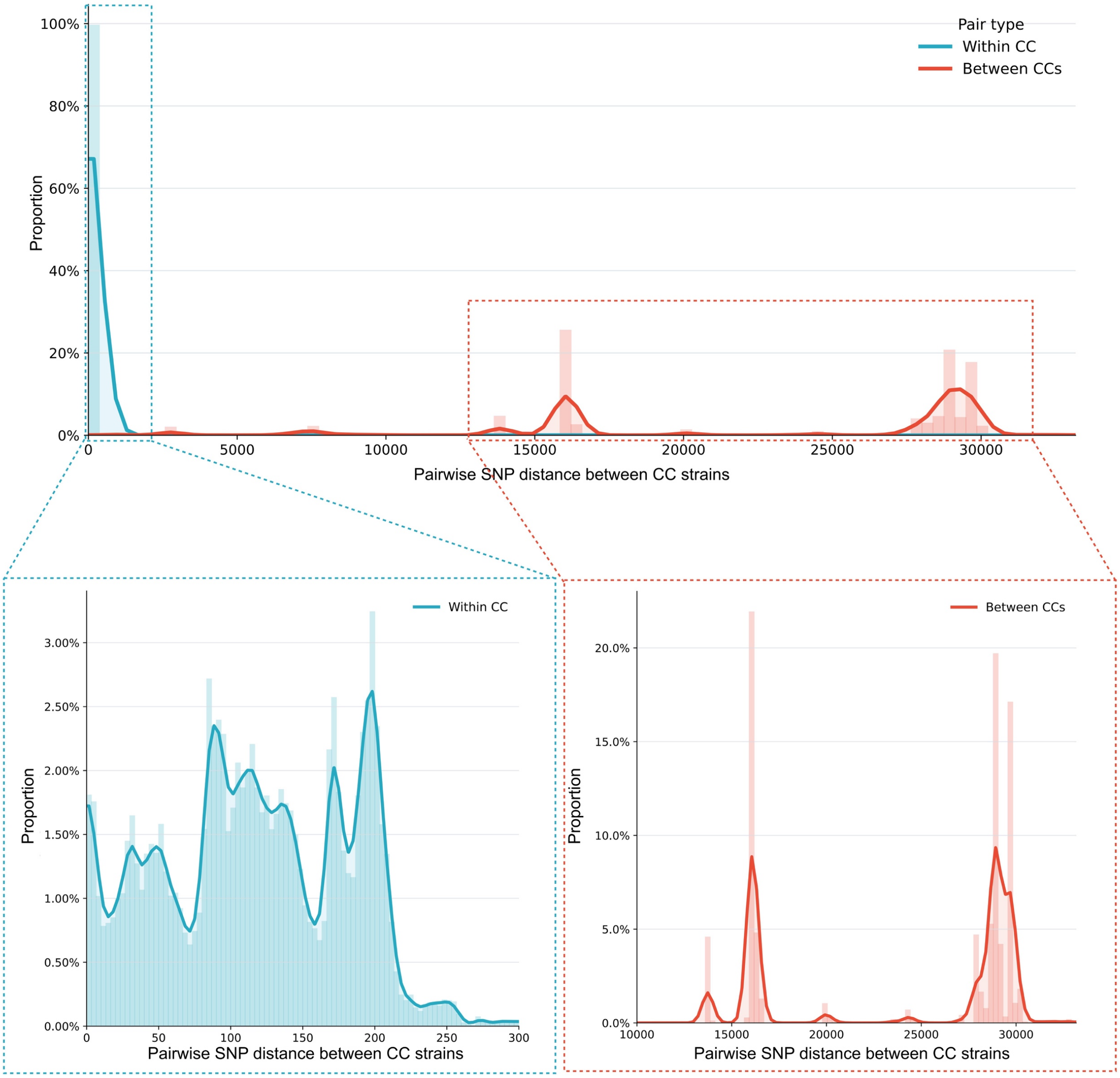


**Supplementary Figure 9. Discontinuous distribution of pairwise SNP distances among MAB strains.** Distribution of pairwise SNP distances between strains belonging to the same circulating clone (within CC) and strains belonging to different circulating clones (between CCs). The lower panels show enlarged views of the within-CC and between-CC distance ranges. The scarcity of strain pairs with intermediate SNP distances highlights the discontinuous population structure of MAB, characterized by close genetic relatedness within circulating clones and deep divergence between clones.

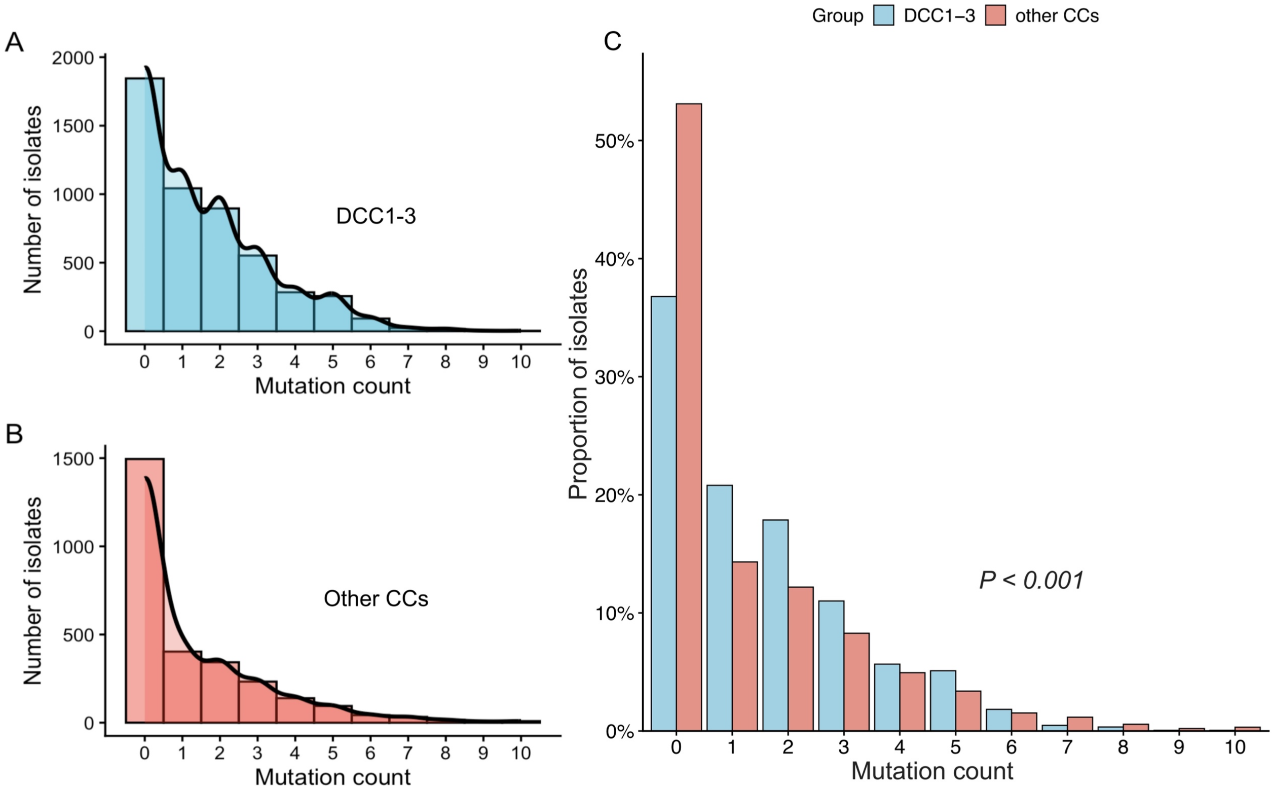


**Supplementary Figure 10. Distribution of nonsynonymous mutation counts per isolate**. (**A**) Histogram with kernel density estimate showing the distribution of nonsynonymous mutation counts among isolates from DCC1–3. (**B**) Corresponding distribution among isolates from the other circulating clones. (**C**) Comparison of the proportion of isolates carrying each number of nonsynonymous mutations in DCC1–3 versus the other circulating clones. The y-axis indicates the proportion of isolates within each group. Differences between groups were assessed using a Wilcoxon rank-sum test ($P < 0.001$).
